# NNMT Loss Drives Cancer Progression by enhancing SAM availability for mTORC1 Signaling and Chromatin Methylation

**DOI:** 10.64898/2026.06.30.734970

**Authors:** Qu Deng, Erick Mitchell-Velásquez, Sharan Venkatesh, Rahul Mannan, Hyein Cho, Mohammed Alhusayan, Aqsa Yashfeen, Ramakrishnan Natesan, Natarajan V Bhanu, Radha Paturu, Javed Siddique, Rohit Mehra, Sooryanarayana Varambally, Benjamin Garcia, David Feldser, Priti Lal, Arul M. Chinnaiyan, Irfan A. Asangani

**Author notes:** Department of Experimental Therapeutics, University of Texas MD Anderson Cancer Center, Houston, TX, USA. Co-first author (Qu Deng, Erick Mitchell-Velásquez).

## Abstract

Aberrant epigenetic reprogramming together with dysregulated mTOR signaling are hallmarks of cancer, where altered chromatin methylation and nutrient-sensing pathways cooperate to drive tumor progression. *S*-adenosylmethionine (SAM), the universal methyl donor, is essential for these processes, yet how tumors sustain elevated SAM availability to support oncogenic transmethylation reactions remains poorly defined. Here, using prostate cancer (PCa) as a model system, we identify nicotinamide N-methyltransferase (NNMT) as a critical metabolic-epigenetic regulator and tumor suppressor. Using a prostate-specific *Nnmt* knockout mouse model, we demonstrate that NNMT loss accelerates PCa progression, particularly in the context of *Pten* deletion, resulting in infiltrating carcinoma and reduced survival. Mechanistically, NNMT functions as a “SAM-sink,” and its loss increases intracellular SAM abundance, thereby activating mTORC1 signaling through SAMTOR-dependent sensing and broadly enhancing chromatin methylation. In human PCa, recurrent genomic deletions of NNMT occur in up to 7% of cases, and NNMT protein expression is largely absent in primary tumors and metastases. NNMT-deficient PCa cells exhibit elevated SAM:SAH ratios, increased histone methylation, and heightened mTORC1 activity, enabling sustained tumor growth even under dietary methionine-restriction (MR). Notably, combined MR and pharmacologic mTORC1 inhibition synergistically suppresses the growth of NNMT-deficient tumors, revealing a previously unrecognized therapeutic vulnerability. Collectively, these findings establish NNMT as a key tumor suppressor that constrains SAM-driven epigenetic and signaling programs in PCa and suggest a rational, diet-based therapeutic strategy for advanced cancers with NNMT loss.

## Introduction

Changes in the methylation status of proteins, nucleic acids, lipids, and metabolites are fundamental in cancer development^1^. In particular, aberrant methylation of histones and DNA reshapes the epigenetic landscape of tumor cells, altering gene expression programs that drive malignant progression. These methylation reactions depend on cellular metabolism, linking nutrient availability to chromatin state and transcriptional regulation. A central metabolite in this process is *S*-adenosylmethionine (SAM), the universal methyl donor for nearly all methyltransferase reactions. Consequently, intracellular SAM abundance and the SAM:SAH ratio are critical determinants of cellular methylation potential and epigenetic regulation^2–6^.

Prostate cancer (PCa) displays extensive epigenetic remodeling and metabolic rewiring that support tumor initiation, progression, and therapy resistance^7–9^. Nutrient availability influences the abundance of metabolites critical for epigenetic modifications, while genetic mutations and epigenetic remodeling enable cancer cells to adapt and exploit these metabolic pathways to biosynthesis and growth^10,11^.Histone methyltransferases (HMTs) and DNA methyltransferases (DNMTs) generate chromatin states associated with transcriptional activation or repression, and dysregulation of these methylation marks is strongly linked to aggressive disease and poor clinical outcomes in PCa^12–24^. These processes require a sustained supply of SAM, which also fuels other transmethylation reactions important for prostate tumor biology, including polyamine biosynthesis^25–27^. Indeed, dysregulated SAM metabolism and elevated SAM:SAH ratios have been observed in advanced PCa^26,28^. However, the mechanisms by which tumors maintain elevated SAM availability to support oncogenic transmethylation reactions remain poorly understood.

Beyond its role in chromatin methylation, SAM also functions as a metabolic signaling molecule. Cellular SAM levels are monitored by SAMTOR, a nutrient sensor that regulates mTORC1 signaling, thereby linking one-carbon metabolism to cellular growth control^29^. Thus, mechanisms that alter SAM abundance can simultaneously influence epigenetic regulation and mTORC1-dependent metabolic programs. Understanding how tumors maintain SAM levels may therefore reveal mechanisms that coordinate chromatin regulation with nutrient-sensing pathways. The concentration of SAM in cells is primarily regulated by the one-carbon metabolism, a process that involves the entry of amino acids into the one-carbon cycle^6,30^. Within this cycle, methionine is converted into SAM through the enzyme MAT2A. Recent studies have revealed that nicotinamide (NAM) metabolism significantly influences cellular SAM pools^31,32^. The enzyme nicotinamide N-methyltransferase (NNMT) utilizes SAM to convert nicotinamide into metabolically inert 1-methylnicotinamide (1-MNAM) and *S*-adenosylhomocysteine (SAH)^33–35^ (**Fig.1A**). Through this reaction, NNMT acts as a metabolic “SAM sink,” diverting SAM from chromatin methylation reactions and thereby influencing epigenetic states. Consistent with this concept, NNMT dysregulation has been reported across multiple cancer types, where its effects on tumor progression appear highly context dependent. In several cancers and cancer-associated fibroblasts, elevated NNMT expression depletes intracellular SAM pools, reducing histone and DNA methylation and promoting tumor progression^31,32,36,37^. In contrast, the role of NNMT in prostate cancer remains poorly defined. While a few reports have described elevated NNMT expression in select PCa models^38,39^, a comprehensive understanding of NNMT status and its functional consequences for metabolism, chromatin regulation, and tumor progression in PCa is lacking. Given that PCa exhibits extensive chromatin methylation and high demand for transmethylation reactions, we hypothesized that tumor-specific loss of NNMT could sustain SAM availability, thereby enhancing epigenetic methylation and growth-promoting signaling pathways. Therefore, in this study, we generated a prostate-specific *Nnmt-*null mouse model to explore its role in PCa. Loss of NNMT in the *Pten*-null background accelerated infiltrating carcinoma progression, significantly increasing mortality. Single-cell transcriptome analysis identified mTORC1 signaling as a key pathway in the *Nnmt* and *Pten* double knockout, and dietary methionine restriction revealed a critical role of mTORC1 signaling *via* SAMTOR sensing of SAM in promoting growth. Importantly, in human PCa, focal genomic deletions at the *Nnmt* locus were observed in up to 7% of cases, with NNMT protein expression largely absent in primary and metastatic tumors. Manipulating NNMT expression in PCa cell lines demonstrated reciprocal changes in the SAM:SAH ratio, resulting in global DNA methylation alterations, changes in euchromatin and heterochromatin marks, mTORC1 activity, and tumor growth. Importantly, dietary methionine restriction in the NNMT-negative PCa xenograft model created a predictable vulnerability that could be effectively targeted with mTORC1 inhibitors, offering a potential therapeutic strategy for advanced PCa with NNMT loss. Together, these findings identify NNMT as a key regulator of SAM metabolism that constrains epigenetic and metabolic signaling programs driving prostate cancer progression.

**Figure 1.**
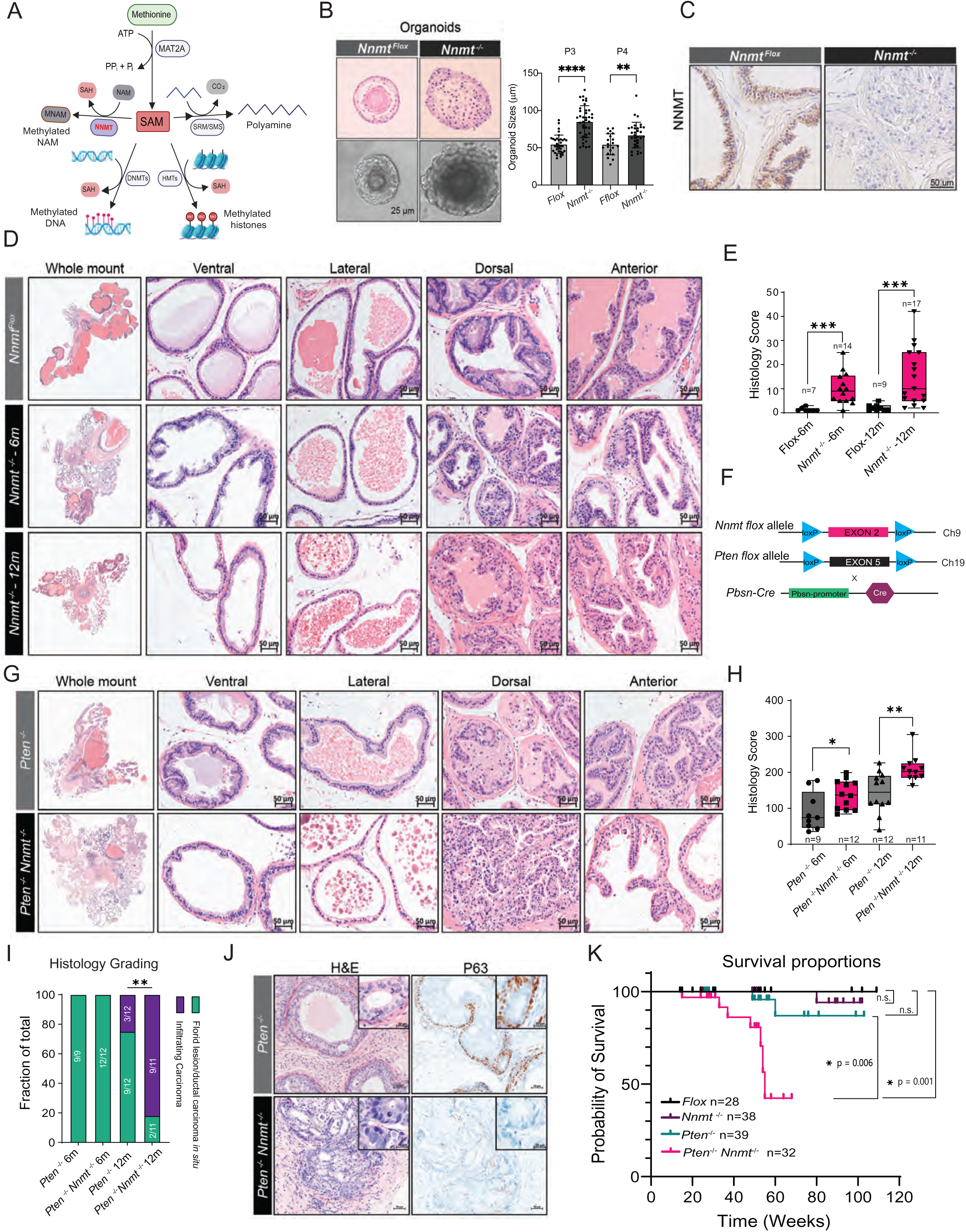
*Nnmt* deletion promotes prostate cancer progression in mice. (A) Schematic of critical SAM-utilizing cellular reactions. (B) *Nnm*t-null prostate organoids exhibit increased proliferation. Organoids formed by control and *Nnmt* knockout mouse prostate epithelial cells. *Left,* Representative hematoxylin and eosin (H&E) and widefield images of organoids formed by the indicated prostate epithelial cells. *Right,* Cre-mediated deletion of flox allele via adeno-Cre transduction during culture. P3 and P4 indicate passages. (C) Immunohistochemistry (IHC) showing NNMT expression in flox control and *Pbsn-Cre Nnmt^-/-^*prostate. (D) Prostate-specific *Nnmt* deletion results in epithelial hyperplasia (Grade I lesion) - H&E images. (E) Histology scores of flox control and *Pbsn-Cre Nnmt^-/-^* prostate at 6 and 12 months. (F) Construction and breeding strategy of prostate-specific *Nnmt* and *Pten* double knockout mouse model. (G-H) *Nnmt* deletion combined with *Pten* loss results in infiltrating carcinoma in the prostate. Histology scores of *Pten^-/-^* and *Pten^-/-^Nnmt^-/-^* prostates at 6 and 12 months. H&E images of *Pten^-/-^*and *Pten^-/-^Nnmt^-/-^* prostate at 12 months. (I) Fractions of total histology grades of *Pten^-/-^* and *Pten^-/-^Nnmt^-/-^*prostates at 6 and 12 months, highlighting high prevalence of infiltrating ductal carcinoma in *Pten^-/-^Nnmt^-/-^*. Grades: III (florid atypical intraductal carcinoma) and IV (infiltrating ductal carcinoma). (J) Representative IHC images of P63 staining showing infiltrating carcinoma developed in *Pten^-/-^Nnmt^-/-^* prostate at 12 months. (K) Kaplan-Meier survival curve of flox control, *Nnmt^-/-^*, *Pten^-/-^,* and *Pten^-/-^Nnmt^-/-^* mice. Statistical significance was assessed using Log-rank (Mantel-Cox) test. Statistical significance for panels B, E, H, and I was assessed with unpaired two-sided Student’s t-tests (*P < 0.05, **P < 0.01, ***P < 0.001 ****P < 0.0001)

## Results

### *Nnmt* deletion promotes PCa progression in a genetically engineered mouse model

The role of tumor cell intrinsic NNMT remains poorly understood due to the reliance on cell line-based studies; therefore, we hypothesized that a conditional Nnmt knockout mouse model would enable physiologically relevant, long-term in vivo analyses across the continuum of tumor development, and accordingly used CRISPR-Cas9 technology to engineer mice in which *Nnmt* exon 2 is flanked by loxP sites (*Nnmt^flox^* mice) (**Fig. S1A**). First, we assessed the regenerative ability of *Nnmt^-/-^* prostate epithelium using organoid cultures following Cre adenovirus-mediated *Nnmt* deletion *in vitro* (**Fig. S1B-D**)^40^. Interestingly, *Nnmt^-/-^* organoids exhibited sustained growth, forming larger and denser structures compared to wildtype controls, and H&E staining revealed their less differentiated and hyperplastic phenotypes, in contrast to cystic structures of wildtype organoids recapitulating normal prostate gland architecture (**Fig. 1B and Fig. S1E**). Next, we generated prostate-specific *Nnmt* knockout mice by crossing the *Nnmt^flox^* with probasin Cre (*Pb-Cre)* mice^41^. Immunohistochemistry (IHC) confirmed nearly complete loss of NNMT protein in the luminal epithelium of the prostate in the *Nnmt^-/-^* mice (**Fig. 1C** and **Fig. S1F**). Although no cancerous lesions were evident, histopathological analysis revealed significant epithelial hyperplasia (Grade I lesion), particularly in the dorsal lobes, in 6- and 12-month-old *Nnmt^-/-^* mice (**Fig. 1D, 1E** and **Fig. S1H**). These findings underscore the importance of NNMT activity in maintaining prostate epithelial differentiation and suggest that its loss may promote disease progression when combined with other critical drivers of the disease.

PTEN is a well-established tumor suppressor in PCa, and conditional *Pten* loss in the mouse has been reported to show neoplasia at a later stage of life^42^. In humans, *Pten* deletions and mutations are relatively low in primary early phases of PCa^43^ but become increasingly more frequent in advanced metastatic disease^44,45^. Given these observations, we hypothesized that PTEN deletion might synergize with NNMT loss to drive aggressive PCa progression in our genetically engineered mouse (GEM) model. To test this, we crossed *Nnmt^flox^* mice with *Pb-Cre* and *Pten ^flox^* line to generate prostate-specific *Pten^-/-^Nnmt^-/-^* mice (**Fig. 1F** and **Fig. S1G**). At six-months, *Pten^-/-^Nnmt^-/-^*mice exhibited significantly increased genitourinary size/weight and higher histology scores compared to *Pten^-/-^* mice (**Fig. 1G** and **Fig. S1I-J**). By 12 months, histopathological analysis showed substantial Grade IV infiltrating carcinoma in the dorsal lobes of *Pten^-/-^Nnmt^-/-^* mice (**Fig. 1H**). Specifically, 9 out of the 11 *Pten^-/-^Nnmt^-/-^* mice developed infiltrating carcinoma affecting up to 20% of the prostate, compared to only 3 out of 12 *Pten^-/-^* mice, where lesions affected just 8% of the prostate (**Fig. 1I**). IHC analysis further revealed defining features of infiltrating adenocarcinoma in the *Pten^-/-^Nnmt^-/-^*prostates, including loss of basal cells marked by P63, disrupted basal lamina marked by Smooth Muscle Actin (SMA), retention of androgen receptor (AR) expression, and high mitotic activity (**Fig. 1J** and **Fig. S1K-L**). These differences were associated with significantly reduced survival; greater than 50% of *Pten^-/-^Nnmt^-/-^* mice succumbed to the disease by week 55, primarily due to bladder obstruction, renal failure, or other potential systemic causes (**Fig. 1K**). Collectively, these findings demonstrate that NNMT loss drives prostate cancer progression in the context of *Pten* deficiency, establishing the *Pten^-/-^Nnmt^-/-^* model as strong evidence for the role of NNMT in promoting PCa progression *in vivo*.

### *Nnmt* deletion is associated with increased mTORC1 signaling

The expression of NNMT has been associated with changes in histone methylation states, including H3K27me3 and H3K36me3^36,46^, which are linked to transcriptional silencing and activation, respectively. To investigate the connection between NNMT loss and the epigenetic and transcriptional alterations driving disease progression *in vivo* (Fig. 1), we evaluated the levels of these histone marks in *Pten^-/-^Nnmt^-/-^* and *Pten^-/-^* prostates from age-matched 10–12-month-old mice. IHC analysis revealed a significant increase in the levels of H3K27me3 and H3K36me3 in lesions from *Pten^-/-^Nnmt^-/-^* prostates compared to *Pten^-/-^* controls (**Fig. 2A**). This suggests an impact of NNMT loss on these specific histone methylation marks *in vivo*, potentially correlating with altered transcriptional state. To investigate transcriptional changes and associated molecular pathway alterations caused by NNMT loss, we performed single-cell RNA sequencing (**Fig. 2B** and **Fig. S2A**). The analysis revealed a marked shift in the epithelial population, with an increased luminal-to-basal ratio observed in *Pten^-/-^Nnmt^-/-^* prostates compared to *Pten^-/-^* controls (**Fig. 2B** and **Fig. S2B**). Gene set enrichment analysis (GSEA) identified mTORC1 signaling as the most enriched pathway within the luminal populations of *Pten^-/-^Nnmt^-/-^*prostate, accompanied by significant enrichment in metabolic pathways including adipogenesis, fatty acid metabolism, and androgen response (**Fig. 2D, 2E**, and **Fig. S2C**). These findings highlight the impact of NNMT loss in driving metabolic and transcriptional reprogramming. Consistent with this, elevated mTORC1 activity was further validated by IHC analysis of downstream effectors, including phospho-S6 ribosomal protein and phospho-4E-BP1 (Thr37/46), both of which were markedly increased in *Pten^-/-^Nnmt^-/-^* prostate tissue (**Fig. 2F**).

**Figure 2.**
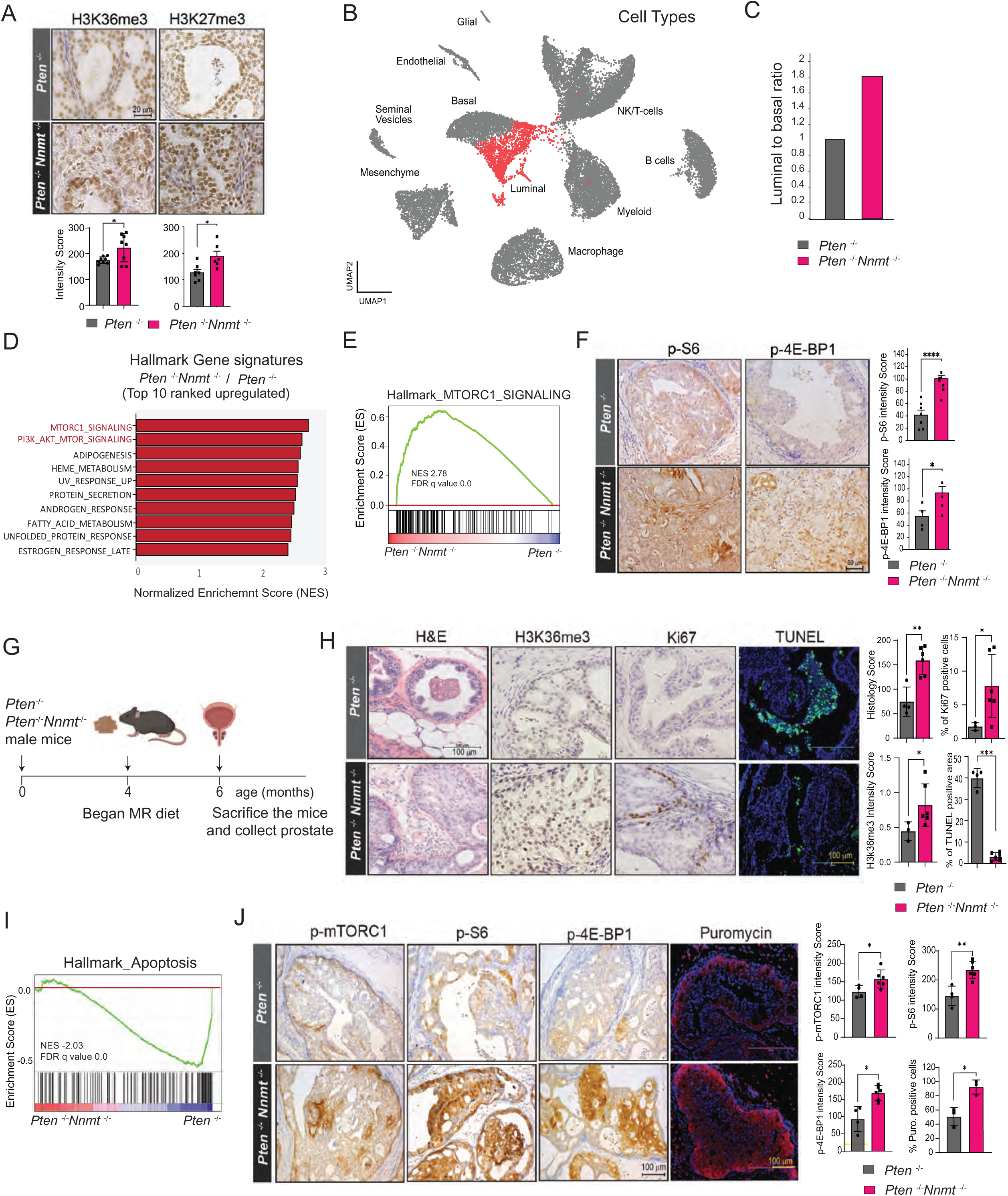
NNMT loss activates mTOR and reduces sensitivity to dietary methionine restriction. (A) *Nnmt* deletion is associated with increased histone methylation. *Top:* Representative IHC images of H3K36me3 and H3K27me3 in *Pten^-/-^Nnmt^-/-^* and *Pten^-/-^* prostates at 12 months. *Bottom*: Quantification. (B) UMAP of scRNA-seq showing cell populations in *Pten^-/-^Nnmt^-/-^* and *Pten^-/-^* prostates (n =3 mice each), highlighting luminal epithelial cells. (C) Luminal to basal cell ratio is shown for each condition. (D) Top 10 enriched Hallmark gene signatures from GSEA of *Pten*^-/-^*Nnmt*^-/-^ and *Pten*^-/-^ prostate epithelial cells. (E) GSEA plot for the Hallmark mTORC1 signaling genes comparing *Pten*^-/-^*Nnmt*^-/-^ vs *Pten*^-/-^ prostate luminal epithelial cells. (F) Representative IHC images and intensity score of phospho-S6 and phospho-4E-BP1 in *Pten^-/-^Nnmt^-/-^*and *Pten^-/-^* prostate at 12 months. (G) Schematic of dietary methionine restriction (MR) experiment in the *Nnmt-*null prostate GEM model. (H) H&E, H3K36me3, Ki67 staining, and TUNEL assay in *Pten*^-/-^ and *Pten*^-/-^*Nnmt*^-/-^ prostate from the MR diet-fed mice. Quantified histology scores, IHC, and TUNEL staining are shown. (I) Bulk RNA-seq derived GSEA plot for Hallmark apoptosis genes in *Pten^-/-^Nnmt^-/-^* vs *Pten^-/-^* prostate tissue (n = 2, biological replicates). (J) Representative IHC and IF images of mTORC1 pathway proteins and puromycin staining (*in vivo* SUnSET assay), respectively. Quantifications shown. Statistical significance was determined using unpaired two-sided Student’s t-tests (*P < 0.05, **P < 0.01, ***P < 0.001 ****P < 0.0001)

Building on these observations, we speculated that NNMT loss modulates mTORC1 activity by altering SAM metabolism. As NNMT functions as a major “SAM-sink,” its loss may result in elevated intracellular SAM levels, potentially activating mTORC1 signaling through SAM sensing by SAMTOR^29,47–49^. We further reasoned that NNMT-deficient PCa, due to increased SAM availability, might exhibit resistance to reduced methionine intake – a dietary approach proposed to be beneficial in cancer management^50^. To test this, we placed four-month-old *Pten^-/-^* and *Pten^-/-^Nnmt^-/-^* mice on a methionine-restricted (MR) diet (0.12% of methionine, w/w) for two months (**Fig. 2G**). Under MR conditions, marked histological differences emerged between *Pten^-/-^* and *Pten^-/-^Nnmt^-/-^* prostates (**Fig. 2H**). The *Pten^-/-^* prostates responded to the MR-diet, with most glands shrinking and maintaining benign epithelium, characterized by negligible mitotic activity (Ki-67) and increased apoptosis (TUNEL positivity). In contrast, *Pten^-/-^Nnmt^-/-^*prostates were resistant to the MR-diet, displaying highly cellular prostatic glands, increased mitotic activity, no apoptosis, and elevated H3K36me3 levels (**Fig. 2H**). To uncover the molecular mechanism underlying these differential responses, we performed bulk RNA sequencing of the entire prostate (**Fig. S2D**). The analysis revealed significant enrichment of multiple Hallmark gene signatures in the *Pten^-/-^Nnmt^-/-^* condition, including AR and mTOR-associated pathways (**Fig. S2E**). Conversely, pathways associated with apoptosis, inflammation, and p53 were among the top downregulated in *Pten^-/-^Nnmt^-/-^*prostates (**Fig. 2I** and **Fig. S2E)**. Importantly, prominently enriched signatures, including fatty acid metabolism, AR signaling, protein secretion, and unfolded protein response, are known to intricately connected to mTORC1 signaling^51,52^. Numerous genes within these pathways are under the direct transcriptional control by factors such as SREBP1 and MYC, both of which are themselves positively regulated by mTORC1^53,54^. Since mTORC1 signaling is the master regulator of the metabolism and cell growth by sensing nutrient and signaling inputs^51^, we examined phospho-mTORC1 and its downstream targets, including phospho-S6 ribosomal protein and phospho-4E-BP1. The IHC analysis revealed a significant increase in their levels in *Pten^-/-^Nnmt^-/-^* condition under the MR-diet (**Fig. 2J**). To assess the functional consequences of increased mTORC1 activity, we conduced puromycin-incorporation assay (SUnSET assay)^55^ to evaluate the rate of translation *in vivo*. This assay demonstrated a marked increase in *de novo* protein synthesis in *Pten^-/-^Nnmt^-/-^* prostates (**Fig. 2J**). These findings strongly suggest that NNMT-deficient PCa depends on mTORC1 signaling to sustain key transcriptional and metabolic pathways, enabling tumor growth despite low-methionine conditions - a characteristic feature of the nutrient-deprived, hypoxic tumor microenvironment driven by uncontrolled proliferationg^1,11^.

### NNMT modulates mTORC1 signaling through SAMTOR sensing of SAM

mTORC1 functions as a master regulator of cell growth and homeostasis, responding to diverse environmental cues, including methionine through SAM sensing^29,49^. SAM regulates mTORC1 activity by disrupting its negative regulator, the SAMTOR-GATOR complex, binding directly to SAMTOR with a dissociation constant of approximately 7µM^29^. Methionine deprivation typically reduces SAM levels below this dissociation constant, enabling the association of SAMTOR and GATOR1 complex, thereby inhibiting mTORC1 signaling in a SAMTOR-dependent manner ^29,48^. Hence, we posited that NNMT, through its “SAM-sink” activity, depletes the cellular SAM pool below this threshold, facilitating the formation of SAMTOR and GATOR1 complex and consequently inhibiting mTORC1 signaling (**Fig. 3A**)^29,47^. To test this, we first overexpressed NNMT in mouse primary prostate organoids and evaluated mTORC1 signaling (**Fig. S3A**). NNMT overexpression resulted in decreased phospho-mTORC1 and phospho-S6K levels compared to controls (**Fig. 3B** and **3C**). Next, we examined the interaction between SAMTOR and DEPDC5, a component of the GATOR1 complex, in HEK-293T cells cultured with or without methionine and with or without NNMT expression. As anticipated, methionine depletion enhanced the interaction between SAMTOR and DEPDC5. Importantly, NNMT expression further strengthened their interaction, even under methionine replete conditions (**Fig. 3D**). This enhanced interaction correlated with reduced phosphorylation of ectopically expressed S6 kinase and a decrease in global protein synthesis, as assessed by puromycin incorporation assay, serving as direct indicators of attenuated mTORC1 activity (**Fig. 3E** and **3F**). These findings were further corroborated in human PCa - VCaP cells overexpressing NNMT, which exhibited decreased levels of phospho-mTORC1, phospho-S6 Kinase, and protein synthesis, and elevated phospho-AMPK to switch off anabolic biosynthesis^56^, compared to the parental controls under physiological methionine conditions (**Fig. 3G, 3H,** and **Fig. 3SB**). These data suggest that NNMT, through its “SAM-sink” activity, coordinates with SAMTOR to directly influence mTORC1 signaling.

**Figure 3.**
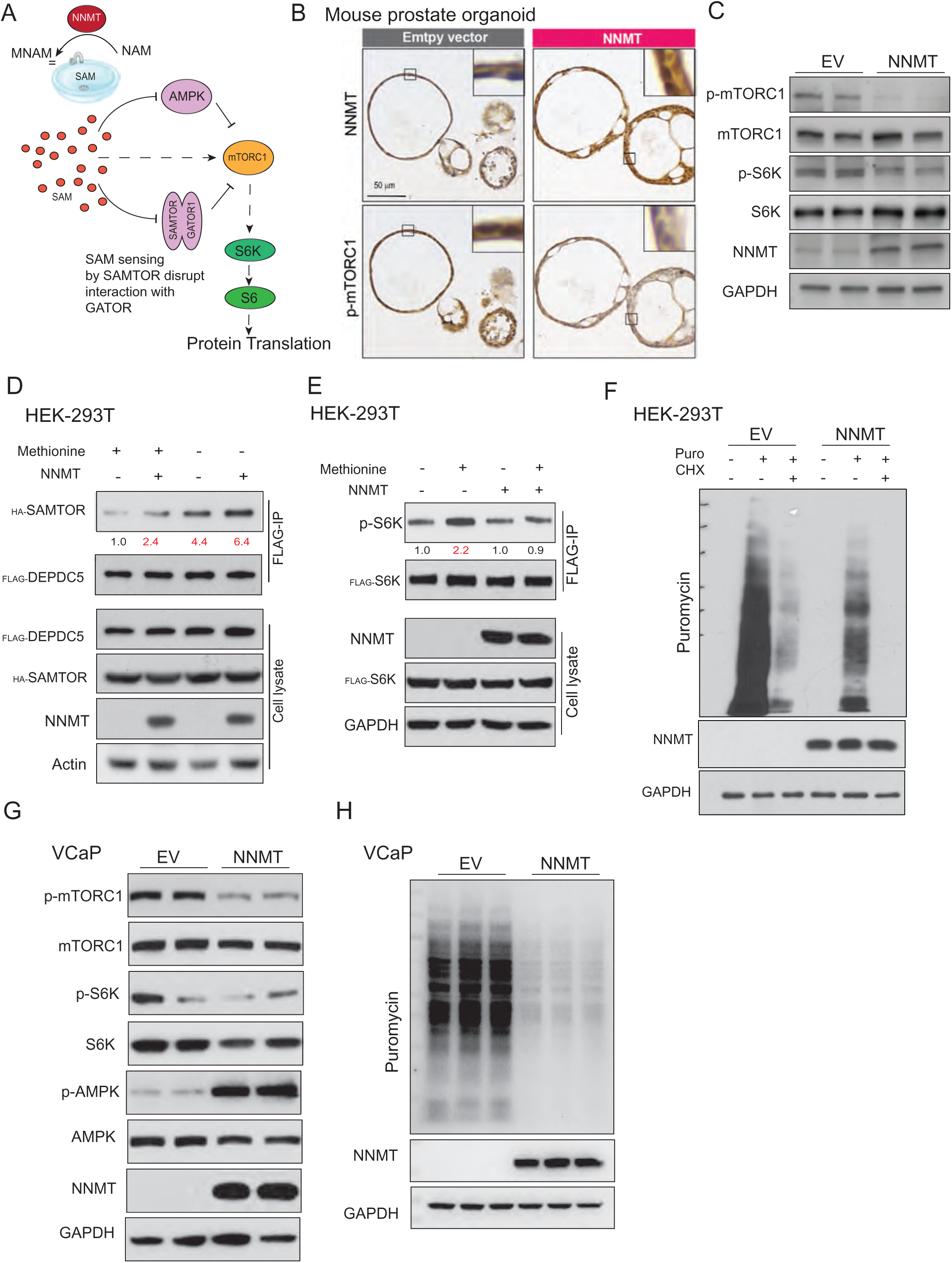
NNMT modulates mTORC1 signaling through SAMTOR sensing of SAM. (A) Schematic model of NNMT as a SAM sink regulating mTORC1 signaling. (B) NNMT overexpression downregulates mTORC1 signaling in mouse prostate organoids. Representative IHC images of NNMT and p-mTORC1. (C) Immunoblot showing reduced mTORC1 signaling in NNMT-overexpressing mouse prostate organoids. GAPDH was used as a loading control. (D) NNMT overexpression increases SAMTOR-DEPDC5 interaction. FLAG immunoprecipitates from HEK-293T cells stably expressing NNMT or vector control with transient transfection of FLAG-DEPDC5 cDNA along with hemagglutinin (HA)-tagged SAMTOR cDNA for 48hrs, followed by 4hrs culture in complete media containing 100μM methionine or methionine deprived media. FLAG immunoprecipitates and cell lysates were analyzed by immunoblotting for the indicated proteins. Numbers below HA-SAMTOR blot indicate normalized values. (E) Attenuated S6K phosphorylation in NNMT-overexpressing cells stimulated with methionine. FLAG immunoprecipitates were prepared from HEK-293T cells stably expressing NNMT or vector control with FLAG-S6K cDNA. Cells were grown in methionine deprived media for 16hrs and stimulated with 100μM methionine for 4 hrs. FLAG immunoprecipitates and cell lysates were analyzed by immunoblotting for the indicated proteins. Numbers below p-S6K blot indicate normalized values. (F) Decreased protein translation in NNMT-overexpressing cells measured by SUnSET assay. Immunoblots showing puromycin-labelled proteins and NNMT overexpression in HEK-293T cells. Cells were grown in regular media and were chased with 10μg/ml puromycin (Puro) for 10 mins before harvesting for protein lysis. 20μg/ml of cycloheximide (CHX) was used as a positive control for translation inhibition. (G) NNMT overexpression reduces p-mTORC1/p-S6K and increases p-AMPK in human prostate cancer cells. VCaP cells stably expressing NNMT were grown in physiological methionine concentration (20μM) for 4 hours, and total lysates were analyzed by immunoblotting for the indicated proteins. (H) VCaP cells were grown as in F and incubated with puromycin for 30 minutes before extracting the total proteins to examine active protein translation.

### Recurrent *Nnmt* genomic deletion and silencing in human prostate cancer

Building on our *in vivo* GEM model findings and *in vitro* mechanistic studies, we next examined the genomic status of NNMT in human PCa, together with other key methyltransferases and enzymes involved in polyamine synthesis, which are known to consume high levels of SAM for transmethylation reactions. Although these enzymes exhibited genomic alteration, including amplifications, mutations, deletions, and structural variants, such changes were detected in only a small fraction of both primary and metastatic PCa cases (**Fig. S4A**). In contrast, we identified recurrent homozygous deep focal deletions on chromosome 11, minimally encompassing the *NNMT* and *ZBTB16* genes, in 4–7% of PCa cases (**Fig. 4A** and **4B**). Notably, ZBTB16 (PLZF) has been implicated in prostate tumorigenesis and androgen resistance in preclinical models^57,58^, with its loss previously associated with the upregulation of MAPK signaling^59^. To investigate the expression status of NNMT protein, we performed IHC on tissue microarrays (TMAs), which demonstrated a significant loss of NNMT expression in both primary and metastatic PCa (**Fig. 4C** and **Fig. S4B**). These results were validated in multiple prostatectomy resections and their corresponding metastatic biopsy samples (**Fig. S4C**). NNMT was predominantly expressed in luminal cells of the normal prostate epithelium (**Fig. 4C** and **S4D**). However, its expression was entirely absent in transformed cells in both primary and metastatic tumors. In contrast, stromal and host cells, such as hepatocytes, exhibit strong NNMT positivity in the microenvironment. This suggests that NNMT-negative tumor cells may gain a growth advantage by conserving intracellular SAM, enabling them to adapt to nutrient-deprived environments and sustain proliferation. Interestingly, metabolomic analysis of human PCa samples^28^ revealed significantly enriched SAM levels in advanced metastatic PCa compared to primary tumors and benign prostatic tissues (**Fig. S4E**).

**Figure 4.**
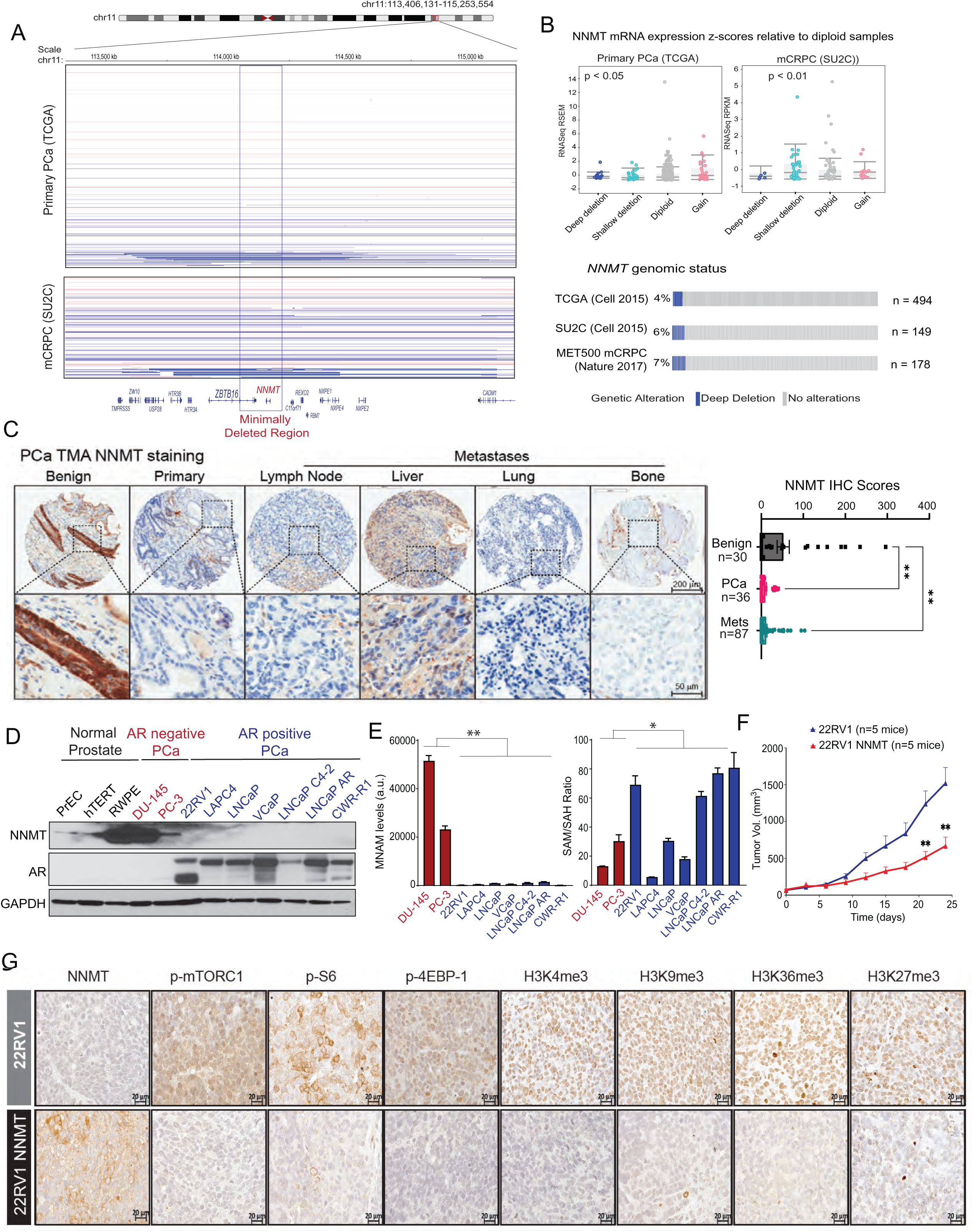
Recurrent *NNMT* genomic deletion and silencing in human prostate cancer. (A) Copy number plot for *NNMT* locus in primary (TCGA) and metastatic (SU2C) prostate cancer datasets. Deep deletion is indicated in blue, and red indicates potential copy gain. (B) *Top:* NNMT mRNA expression correlates with copy number changes in TCGA and SU2C datasets (deep deletion vs diploid). *Bottom:* Oncoprint from CBioPortal highlights deep deletion of the *NNMT* locus in primary and metastatic prostate cancer datasets. (C) Reduced NNMT protein expression in prostate cancer. Representative IHC images and quantified scores from TMA cores. Note that only adjacent hepatocytes are positive, while cancer cells in liver metastases are negative for NNMT. (D) Immunoblot showing NNMT and AR in normal and PCa cells. GAPDH was used as a loading control. (E) LC-MS quantification of 1-MNAM and SAM/SAH ratios in the indicated PCa cells. Note the absence of 1-MNAM in NNMT-negative cells. (F) NNMT overexpression suppresses AR-positive 22RV1 xenograft growth in mice. (G) Reduced mTOR signaling and histone methylation in NNMT-overexpressing 22RV1 xenografts. Representative IHC images of the phospho-proteins in mTORC1 pathway and indicated histone methylation marks in control and NNMT-overexpressing 22RV1 tumor xenografts. Statistical significance was assessed using unpaired two-sided Student’s t-test (*P < 0.05, **P < 0.001)

To further characterize NNMT expression, we assessed a panel of 12 prostate cell lines. All three normal prostate epithelial lines and the two AR-negative PCa cell lines showed positive NNMT expression. However, all seven AR-positive PCa cell lines lacked detectable NNMT protein (**Fig. 4D**). This observation is particularly intriguing, as the vast majority of prostate adenocarcinomas are AR-driven^60^, and the clear negative correlation between NNMT and AR expression strongly suggests that NNMT loss, through genomic deletion and transcriptional silencing, is associated with disease progression. Using liquid chromatography, high-resolution mass spectrometry (LC-HRMS) based metabolite measurements, we observed an accumulation of nicotinamide (NAM) and no detectable 1-MNAM (main product of NNMT enzymatic activity) in the seven NNMT negative PCa cells, whereas high levels of 1-MNAM were evident in DU145 and PC3 cells, both of which expressed NNMT (**Fig. 4E** and **Fig. S4F**). This finding is consistent with the loss of NNMT methyltransferase activity in AR-positive PCa cells. Interestingly, despite no major differences in total SAM levels between the cell lines (likely due to methionine saturation in standard media), NNMT-negative cell lines displayed a significant increase in SAM:SAH ratios, indicative of enhanced methylation potential in NNMT-negative cells. This evidence highlights a potential mechanism by which NNMT loss contributes to PCa progression - by conserving SAM levels and enhancing methylation potential, thereby driving transmethylation reactions that promote chromatin methylation and protein translation through mTORC1 signaling.

### NNMT overexpression suppresses PCa growth *via* decreased mTORC1 activity and histone methylation

To investigate the impact of NNMT overexpression on the growth of AR-positive, NNMT-negative PCa cells, we utilized a retroviral system to overexpress NNMT in LNCaP, 22RV1, and VCaP cells (**Fig. S5A**). NNMT overexpression led to a marked reduction in PCa cell growth, as evidenced by decreased proliferation, colony formation, and sphere formation *in vitro* (**Fig. S5B-D**). To determine whether the growth inhibitory effects of NNMT overexpression were mediated by its metabolic product, 1-MNAM, we treated PCa cells with exogenous 1-MNAM. However, 1-MNAM treatment did not affect cell growth, suggesting that the observed phenotype is not directly attributable to this metabolite (**Fig. S5E**). We next extended these findings to an *in vivo* context using 22RV1 and VCaP xenograft model to assess the effects of NNMT overexpression. The results demonstrated a significant reduction in tumor growth *in vivo* in both models (**Fig. 4F** and **Fig. S5F-H**). IHC analysis of the xenograft tumors further revealed robust NNMT expression, which correlated with suppressed mTORC1 signaling and reduced levels of multiple histone methylation marks (**Fig. 4G and Fig. S5I**). In parallel, transcriptomic analysis of human PCa samples revealed that mTORC1 signaling, along with interconnected metabolic pathways such as fatty acid metabolism, AR signaling, protein secretion, and the unfolded protein response^51,52^, was significantly enriched in NNMT-low compared to NNMT-high tumors and was associated with poor overall survival (**Fig. S6A** and **S6B**). Notably, we also observed increased levels of H3K27me3, in lysates derived from PCa tissues compared to benign tissues (**Fig. S6C**). Furthermore, ChIP-seq analysis of H3K27me3 in matched normal and primary prostate tumor samples^61^ revealed dramatic genome-wide enrichment for this transcription silencing mark in PCa (**Fig. S6D**). Together, these findings suggest that NNMT functions as a tumor suppressor in the context of PCa by modulating intracellular SAM levels that directly affect mTOR activity and histone methylation.

### NNMT modulates cellular SAM levels, affecting chromatin methylation and transcription

To investigate the effect of NNMT on SAM metabolism in human PCa cells (**Fig. 5A**), we generated NNMT knockout models using CRISPR-Cas9 (**Fig. S7A**). Mass spectrometry-based metabolite quantification revealed a significant increase in the SAM:SAH ratio and a loss of 1-MNAM in NNMT-deficient DU145 cells (**Fig. 5B** and **Fig. S7B**). Conversely, ectopic expression of NNMT in LNCaP, VCaP, and 22RV1 cells reduced the SAM:SAH ratio (primarily due to SAH accumulation) and markedly increased 1-MNAM levels through enhanced nicotinamide methylation (**Fig. 5C** and **Fig. S7C**). Consistent with previous report^32^, NNMT overexpression did not result in a substantial decrease in total SAM levels, likely reflecting the supraphysiological methionine concentration (100 µM) in standard culture media, which is approximately fivefold higher than circulating methionine levels in human plasma^62^. Given the pronounced alterations in cellular methylation potential, we next examined the role of NNMT in regulating chromatin methylation. Aberrant hypermethylation of CpG islands in promoters and non-CpG methylation within gene bodies is well-established as a driver of PCa progression, as it adversely affects transcription^16–19^. To assess the consequences of NNMT-mediated SAM depletion on DNA methylation under physiologically relevant conditions, cells were cultured in media containing 20 µM methionine. Under these conditions, NNMT knockout cells exhibited a significant increase in 5-methylcytosine (5mC) levels, whereas NNMT-overexpressing cells showed a marked reduction in 5mC signal (**Fig. 5D**). As NNMT-mediated SAM consumption also impacts histone methylation^32,63^, and histone methylation is metabolically sensitive to cellular methylation potential and extracellular methionine concentration^64^ (**Fig. S7D**), we conducted histone post-translational modification (PTM) analysis by mass spectrometry in NNMT-proficient RWPE cells and NNMT-deficient 22RV1 cells. This analysis revealed a striking increase in the number of histone methylation marks in 22RV1 cells relative to RWPE cells, whereas histone acetylation marks showing equal distribution (**Fig. 5E**). These findings were independently validated by immunoblotting of histone extracts from RWPE, 22RV1, and VCaP cells, confirming elevated histone methylation in NNMT-deficient context (**Fig. 5F**). To directly interrogate the causal relationship between NNMT-mediated SAM availability and histone methylation in an isogenic setting, NNMT-overexpressing or NNMT-depleted cells and their respective controls were cultured under physiological methionine conditions. As expected, NNMT overexpression led to a broad reduction in all tested histone methylation marks, whereas NNMT knockdown produced a reciprocal increase (**Fig. 5G**). Importantly, expression of an enzymatically inactive NNMT mutant (Y20A)^65^ failed to alter histone methylation, confirming that the enzymatic activity of NNMT is required for these effects (**Fig. 5H** and **Fig. S7E-F**).

**Figure 5.**
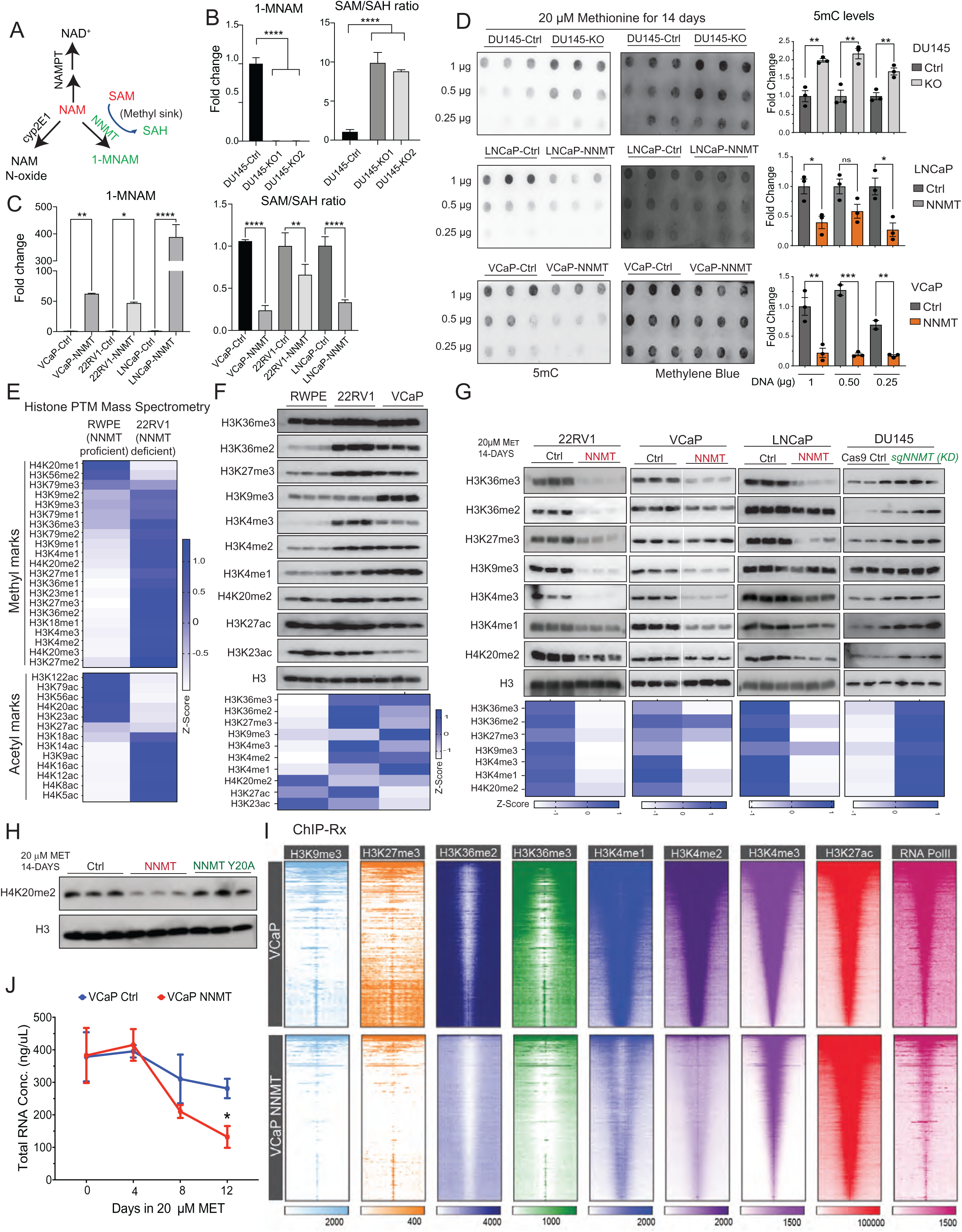
NNMT modulates cellular SAM levels affecting DNA and histone methylation in prostate cancer cells. (A) Schematic of SAM-dependent nicotinamide metabolism by NNMT. (B-C) NNMT depletes cellular SAM levels. Reciprocal changes in NNMT-catalyzed 1-MNAM levels and SAM/SAH ratios were measured by LC-MS in NNMT knockout and overexpressing prostate cancer cells. (D) NNMT expression affects DNA methylation levels. 5mC dot blots and quantification from cells grown in physiological methionine levels. Methylene blue served as the DNA loading control. (E) Effects of NNMT on histone methylation. Z-score heatmap showing histone PTM detected by LC-MS in NNMT-proficient and -deficient cells cultured in regular media (100 μM methionine). (F) Validation of histone PTM changes via immunoblots. Heatmap shows the z-scores of quantified signals normalized to total histone. (G) Immunoblot analysis of various histone methylation marks in NNMT-overexpressing or knockdown cells grown in physiological concentration of methionine. Heatmap shows the z-scores of quantified signals normalized to total histone. (H) Loss of function NNMT Y20A mutant does not affect histone methylation. Immunoblot of H4K20me2 in LNCaP cells overexpressing wildtype NNMT or the NNMT Y20A catalytic-dead mutant. (I) Genome-wide reduction in multiple histone methylation marks detected by ChIP-Rx in NNMT-overexpressing cells. Heatmap depiction of the indicated histone methylation marks, sorted by H3K4me3, in cells grown as in G. H3K27ac, and RNA PolII were used as controls for active transcription marks. (J) NNMT overexpression leads to a decrease in transcription in PCa cells. Total RNA extracted from an equal number of cells grown in physiological concentration of methionine was quantified using TapeStation and plotted (n =3, biological replicates). Statistical significance was assessed using unpaired two-sided Student’s t-test (*P < 0.05, **P < 0.01, ***P < 0.001 ***P < 0.0001, n.s. non-significant).

Although previous studies suggested that specific histone methylation marks, such as H3K4me3^64^, H3K27me3^36,46^, and H3K9me3^63^, are preferentially affected by NNMT expression, our data instead support a substantially broader role for NNMT as a global “SAM sink,” evidenced by a pan-reduction of histone methylation marks upon NNMT overexpression (Fig. 5G). To map the genome-wide distribution of these marks, we performed ChIP-seq for multiple histone methylation marks (H3K4me1, H3K4me2, H3K4me3, H3K36me2, H3K36me3, H3K27me3, and H3K9me3), along with H3K27ac and RNA Polymerase II (RNA PolII), in VCaP cells cultured under physiological methionine conditions for 14 days. Notably, these cells maintained mitotic division, as indicated by the presence of comparable levels of phospho-histone in control and NNMT overexpressing cells (**Fig. S8A** and **S8B**). To enable accurate normalization and quantitative comparison, a defined amount of Drosophila chromatin was included as an internal spike-in control^66^. NNMT expression levels in engineered VCaP cells were comparable to endogenous levels in DU145 cells, effectively ruling out supraphysiological NNMT levels as the underlying cause of the observed phenotype (**Fig. S8A**). ChIP-seq analysis revealed a global reduction in both euchromatin and heterochromatin methylation marks, accompanied by a marked decrease in RNA PolII occupancy in NNMT-overexpressing cells, while H3K27ac levels remained largely unchanged (**Fig. 5I**). Furthermore, analogous patterns of histone methylation loss and diminished RNA Pol II distribution were observed at transcription start sites (TSS) and across open and repressed chromatin genome-wide (**Fig. S8C** and **S8D**). To directly assess the transcriptional consequences of NNMT overexpression, we quantified total RNA levels in a time-course assay. NNMT overexpression in both VCaP and LNCaP cells resulted in a progressive reduction in total RNA content over time (**Fig. 5J** and **Fig. S8E–F**). While we do not anticipate that endogenously NNMT-overexpressing cancer cells experience such dramatic global losses in histone methylation and transcription - likely due to compensatory or adaptive mechanisms - these findings nonetheless underscore the potent SAM-sink activity of NNMT and its capacity to disrupt histone post-translational modification homeostasis. Collectively, these data demonstrate that NNMT depletes intracellular SAM pools, limiting its availability for chromatin methyltransferases, resulting in decreased DNA methylation, histone methylation and overall transcriptional output. These results reveal a profound impact of NNMT activity on the global epigenetic landscape and translational control, including *via* mTOR signaling, highlighting NNMT as a key metabolic–epigenetic regulator capable of reshaping oncogenic transcriptional programs and cancer cell phenotypes.

### NNMT deficiency confers therapeutic vulnerability to mTORC1 inhibitors when combined with methionine-restricted diet

Building on insights from the prostate-specific Nnmt knockout mouse model and complementary mechanistic cell-line studies demonstrating the impact of NNMT “SAM-sink” activity on chromatin methylation and mTORC1 signaling, we hypothesized that methionine restriction (MR) would not impair the growth of NNMT-deficient human PCa due to a compensatory, mTORC1-dependent growth program. This hypothesis was based on the premise that elevated tumor-intrinsic SAM availability in NNMT-deficient cells sustains mTORC1 activation and protein synthesis despite dietary methionine limitation, as observed in *Pten^-/-^Nnmt^-/-^*mouse prostates (**Fig. 2**). To test this, we evaluated the growth of DU145 parental (NNMT-proficient) and isogenic NNMT-deficient xenografts in mice maintained on either control or MR-diet (**Fig. 6A**). Dietary methionine restriction significantly inhibited the growth of NNMT-proficient tumors compared to control diet (**Fig. 6B** and **S9A**). In contrast, NNMT-deficient tumors exhibited comparable growth under both dietary conditions. As expected, NNMT-deficient xenografts displayed an increased SAM:SAH ratio accompanied by depletion of 1-MNAM, irrespective of dietary condition (**Fig. S9A**). Consistent with observations in *Pten^-/-^Nnmt^-/-^* mouse prostate, NNMT-deficient tumors under MR showed elevated mTORC1 signaling, increased protein synthesis, a high proliferation index, and reduced apoptosis relative to NNMT-proficient tumors (**Fig. 6C-D** and **Fig. S9B**). These findings suggest that NNMT-deficient PCa cells activate mTORC1 signaling, potentially through SAMTOR-mediated sensing of intracellular SAM, to drive protein synthesis and sustain tumor growth under MR conditions. Furthermore, while MR depleted H3K27me3 and DNA 5mC levels in NNMT-proficient DU145 tumors, these epigenetic modifications remained unaffected in NNMT-deficient tumors (**Fig. S9C-D**). Similar growth patterns and robust mTORC1 activity were observed in additional independently derived isogenic single-clone NNMT knockout tumor models (**Fig. S9E-G**). To assess therapeutic vulnerability, we combined MR with the mTORC1 inhibitor Everolimus in established NNMT-deficient DU145 xenografts (**Fig. 6E**). This combination resulted in significant tumor growth inhibition or regression, accompanied by reduced mTORC1 signaling, decreased protein synthesis, and diminished mitotic activity compared to MR alone (**Fig. 6F-G** and **Fig. S9H-I**).

**Figure 6.**
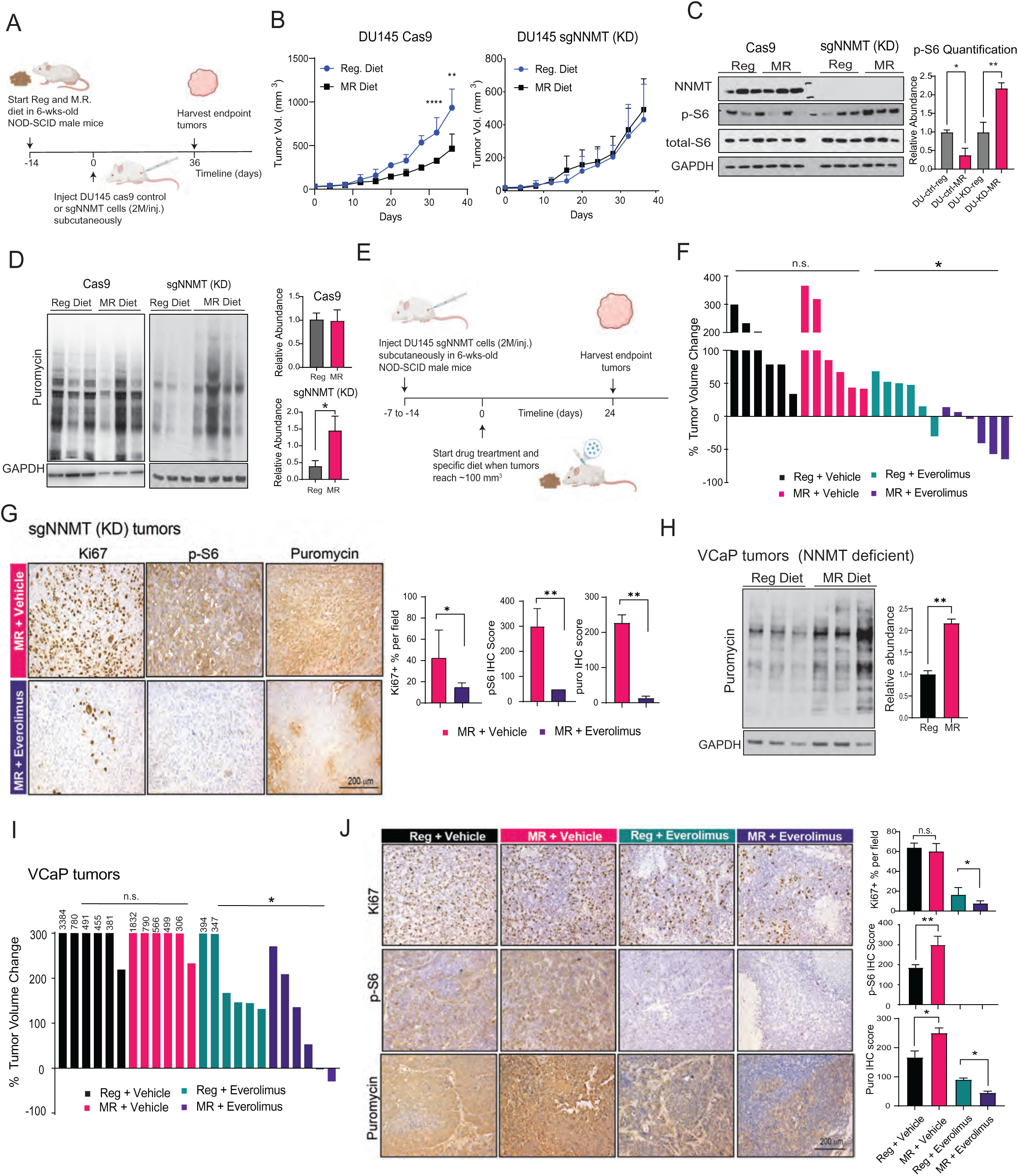
NNMT deficiency creates therapeutic vulnerability to mTOR inhibitors when combined with methionine-restricted diet. (A) Schematic of the experimental design evaluating the effect of a methionine-restricted (MR) diet in isogenic PCa xenograft models with and without NNMT expression. (B) NNMT-knockdown (KD) DU145 xenografts are resistant to MR-diet due to mTOR signaling activation. Mice fed with regular diet or accustomed to MR-diet for two weeks were subcutaneously injected with DU145 Cas9 or NNMT-kd cells. Data are presented as mean tumor volume ± s.e.m. (n = 4-5 mice per group). (C) MR-diet induces mTORC1 signaling in the NNMT-kd tumors, shown by p-S6 levels. *Left:* Immunoblots of tumor lysates*. Right:* Quantification of p-S6 signal intensity. (D) MR-diet increases protein synthesis in NNMT-kd tumors. *Left,* puromycin blots. *Right,* Quantification. Puromycin was injected intraperitoneally in mice 20 min before tumor extraction to assess protein translation rates. (E) Schematic of the experimental design to evaluate MR diet combined with Everolimus as a therapeutic strategy. (F) MR-diet synergizes with Everolimus to reduce tumor growth in NNMT-KD tumors. Waterfall plot shows endpoint tumor volume changes. Everolimus (10mg/kg) was delivered daily through oral gavage. (G) Representative IHC images showing the indicated markers in tumors from MR-diet alone or in combination with Everolimus treated mice. Quantification of the IHC staining is shown. (H) MR-diet increases protein synthesis in NNMT-deficient VCaP tumors. Mice bearing VCaP xenografts were fed on regular diet or MR-diet for 10 weeks. Puromycin i.p. injection in mice was performed 20 min before tumor extraction to monitor the rate of protein translation. Total protein lysate was used for immunoblotting with anti-puromycin antibody. Each lane represents an independent tumor lysate. *Right.* Quantifications. (I) As in F, with NNMT deficient VCaP tumors. (J) MR-diet induced mTOR activation and protein translation are reversed by Everolimus in VCaP tumors. *Left.* Representative IHC images showing the expression of the indicated markers in the four VCaP xenograft experimental groups. *Right.* Quantifications. Statistical significance was assessed using an unpaired two-sided Student’s t-test. *P < 0.05, **P < 0.01, ***P < 0.001 ***P < 0.0001, n.s. non-significant.

We next extended these findings to NNMT-deficient VCaP xenografts under MR. Consistent with NNMT-deficient DU145 model, VCaP tumors were refractory to MR alone and exhibited increased protein translation rates, likely as a compensatory response to methionine limitation (**Fig. 6H** and **S10A**). Similar resistance to MR was observed in NNMT-deficient H460 human lung cancer xenografts, whereas NNMT-proficient A549 xenografts responded to MR, indicating that this NNMT-dependent metabolic adaptation extends beyond prostate cancer. (**Fig. S10B-C**). Notably, combining MR with mTORC1 inhibition in VCaP xenografts produced a strong synergy, resulting in significant tumor growth inhibition than either MR or Everolimus alone (**Fig. 6I** and **Fig. S10D**), accompanied by a more pronounced reduction in protein translation (**Fig. 6J** and **Fig. S10E**). Together, these findings demonstrate that methionine deprivation forces NNMT-deficient tumors to rely heavily on mTORC1 signaling for survival and growth, thereby creating a therapeutically exploitable vulnerability.

## Discussion

SAM occupies a uniquely central position at the intersection of epigenetic regulation and nutrient-responsive growth signaling, serving simultaneously as the universal methyl donor for chromatin modification and as a metabolite sensed by mTORC1 via SAMTOR to control protein synthesis and cellular anabolism. How cancer cells coordinate these dual SAM-dependent outputs - epigenetic remodeling and nutrient-sensing growth signaling – remains incompletely understood. Here, we identify a previously unrecognized role for NNMT as a key metabolic-epigenetic regulator that constrains intracellular SAM availability and thereby coordinates chromatin methylation with nutrient-responsive mTORC1 signaling in PCa. Although NNMT overexpression has been linked to oncogenic reprogramming in multiple tumor types, the biological consequences of NNMT loss in prostate epithelium had remained unexplored. By developing a prostate-specific *Nnmt* knockout mouse model, we demonstrate that NNMT loss accelerates PCa progression to infiltrating carcinoma. Mechanistically, abrogation of nicotinamide methylation leads to SAM accumulation, enhanced chromatin methylation, and sustained mTORC1 activation, collectively promoting oncogenesis within the prostate microenvironment (**Fig. 7**).

**Figure 7.**
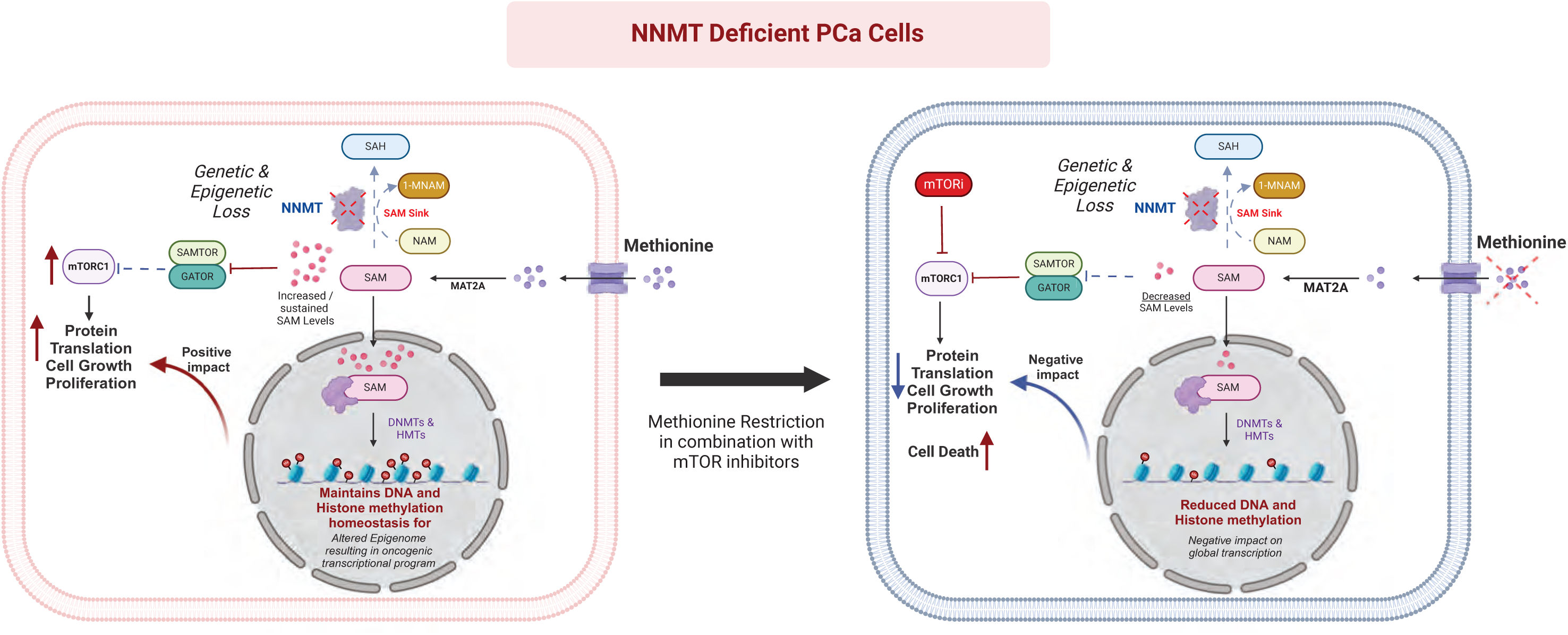
Proposed model illustrating the impact of NNMT deficiency on SAM-dependent epigenetic and metabolic pathways in PCa cells. NNMT deficiency increases SAM availability, which is sensed by SAMTOR, leading to mTOR pathway activation. Excess SAM also supports DNA and histone methylation, altering transcription and gene expression. These effects can be mitigated by a methionine-restricted diet and mTOR inhibitors, ultimately driving the death of NNMT-deficient PCa cells.

Our findings place NNMT within a broader framework of context-dependent cancer metabolism and epigenetic landscape. Elevated NNMT expression correlates with poor prognosis and advanced disease in glioblastoma, gastric cancer, and ovarian cancer, where it promotes tumor progression through SAM depletion and epigenetic reprogramming^31,32,36^. In contrast, our genomic and IHC analyses reveal that NNMT is frequently downregulated or genetically deleted in advanced PCa. We propose that this selective pressure reflects a metabolic advantage in prostate tumors: in a chronically nutrient-limited, hypoxic tumor microenvironment, reducing NNMT expression may conserve methyl-group currency (SAM) and prioritize essential transmethylation reactions and growth signaling rather than diverting SAM into inert 1-MNAM production. In androgen-driven prostate epithelium - where transcriptional output, polyamine metabolism, and chromatin regulation are tightly coupled to proliferation - NNMT loss may therefore enhance fitness by maintaining a high SAM:SAH ratio and supporting methylation-dependent programs under nutrient stress.

This model is supported by our dietary and cross-lineage experiments. NNMT-proficient tumors exhibit sensitivity to MR, consistent with reduced SAM availability limiting methylation-dependent processes. By contrast, NNMT-deficient tumors in both prostate and lung models resist MR monotherapy and instead maintain mTORC1 signaling, consistent with sustained SAMTOR-mediated nutrient sensing despite constrained methionine input. Thus, NNMT status - not tumor lineage per se - emerges as a key determinant of the metabolic response to MR, aligning with a broader principle that tumors optimize SAM allocation according to their dominant constraints (availability of nutrients/oxygen versus demand for methylation and translation).

At a mechanistic level, our data redefine NNMT as a regulator of methyl-group buffering and nutrient sensing. Histones can function as dynamic methyl-group reservoirs, buffering fluctuations in SAM availability in response to metabolic cues ^3,64^. While NNMT overexpression has been proposed to preserve SAM homeostasis under nutrient limitation, our findings demonstrate that NNMT loss disables this buffering “sink”, allowing SAM to remain elevated, thereby promoting global chromatin hypermethylation and enforcing sustained mTORC1-dependent protein synthesis. In this view, NNMT loss does not simply increase methylation potential; it locks tumors into a SAM-high, translation-competent state that is particularly advantageous when exogenous nutrients are limiting, and protein synthesis becomes rate-limiting for growth.

These findings have direct therapeutic implications. Structural and functional studies have established SAMTOR as a molecular switch regulating mTORC1 through the GATOR1–KICSTOR axis^29,48^, and our data suggest that NNMT-deficient tumors exploit this SAM-sensing circuitry to maintain mTORC1 activity under MR. Importantly, this adaptive state creates a precision vulnerability where MR forces NNMT-deficient tumors into heightened reliance on mTORC1, and combined MR with pharmacologic mTORC1 inhibition overcomes resistance and suppresses tumor growth. Early-phase clinical studies indicate that MR is feasible and can modulate systemic one-carbon metabolism in humans, providing a foundation for therapeutic implementation. Short-term MR diets are well tolerated and achievable through defined dietary interventions, including plant-based regimens, and preliminary clinical trials suggest potential synergy with standard therapies^67,68^. These observations provide a practical foundation for NNMT-stratified MR combination trials, in which NNMT expression or genetic status may serve as a biomarker for patient selection. Moreover, because NNMT loss increases methyl-group availability, combining MR and mTOR inhibition with small molecule inhibitors of DNA or histone methyltransferase (e.g., DNMT, EZH2, or NSD2) could further destabilize metabolic-epigenetic adaptability and amplify therapeutic efficacy.

In summary, we identify NNMT as a central metabolic-epigenetic regulator that links its “SAM-sink” activity to chromatin methylation and SAMTOR-mTORC1 signaling. By constraining intracellular SAM availability, NNMT coordinates epigenetic state with nutrient-responsive growth signaling, and its loss enforces a state of sustained mTORC1 dependence. While PCa provides the strongest genetic and clinical evidence for NNMT loss in this study, our cross-lineage dietary experiments, together with emerging evidence from other tumor contexts^69,70^, highlights the broader principle that NNMT is a tunable determinant of SAM allocation for epigenetic and metabolic signaling. Together, our work demonstrates a mechanistic framework and therapeutic rationale in which NNMT status stratifies sensitivity to dietary and pharmacologic interventions, revealing a synthetic vulnerability that can be exploited under methionine restriction to target otherwise adaptive mTORC1-driven tumors, including advanced PCa.

## Acknowledgments

We extend our thanks to the Molecular Pathology and Imaging Core and the Center for Molecular Studies in Digestive and Liver Diseases from UPenn for their assistance with tissue processing and sectioning. We thank the Transgenic & Chimeric Mouse Facility for their expertise in generating genetically engineered mouse model and the Children’s Hospital of Philadelphia Center for Applied Genomics for conducting scRNA-seq. We are also grateful to The Wistar Institute’s Proteomics & Metabolomics Facility for their technical support. Special thanks to Dr. Katherine Wellen for useful discussion, and Gabriel Raytsis for technical assistance. Q.D. is supported by the Abramson Family Cancer Research Institute Postdoctoral Fellowship and the Marlene Shlomchik Fellowship in Cancer Research. Research in Asangani lab is supported by grants from the NIH (5R01CA249210-05, 1R01CA299870-01) and American Cancer Society to I.A.A.

## Author Contributions

I.A.A. and Q.D. conceived the study. Q.D., E.M.V., and I.A.A. designed the experiments. Q.D. established, maintained, and characterized the *Nnmt* and *Pten* knockout transgenic and immunodeficient mouse colonies and performed the *in vivo* experiments with technical assistance from E.M.V. Q.D., R.M., and P.L. performed histopathological evaluations of transgenic mouse models and prostate cancer patient samples. Q.D. established NNMT knockout, overexpression, and mutant transgenic cell line models and performed metabolic profiling, immunoblotting, immunoprecipitation, and puromycin tracing experiments. E.M.V. performed 5-meC dot blotting, histone marker immunoblotting, RNA sequencing, ChIP sequencing, and mouse organoid experiments with assistance from Q.D., H.C., A.Y., and N.V.B. Sh.V., M.A., R.N., and I.A.A. performed bioinformatic analyses. J.S., S.V., D.F., B.G., and A.M.C. provided reagents and technical information. Q.D., E.M.V., and I.A.A. wrote the manuscript. All authors read the manuscript and provided comments. I.A.A. supervised the overall study.

## Data and Software Availability

All sequencing data generated in this study have been deposited in the Gene Expression Omnibus under accession GSE285539 (token: slajkocijderxkb).

---

**Figure S1.**
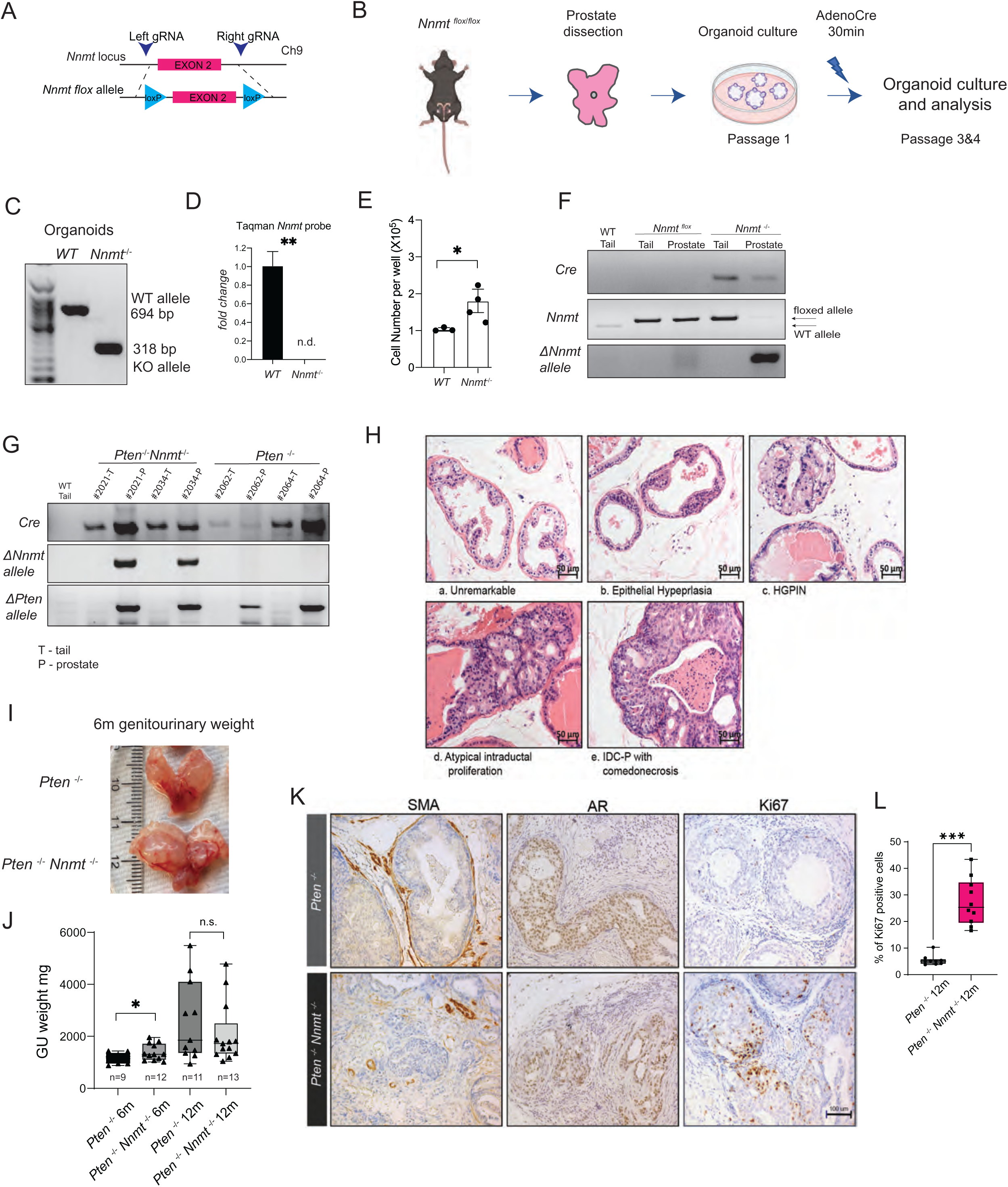
*Nnmt* deletion promotes prostate cancer progression in mice. (A) Construction strategy for CRISPR-Cas9-mediated knock-in of flox sites in *Nnmt* allele. (B) *In vitro* knockout of *Nnmt* in prostate epithelial organoids increases proliferation. Schematic of organoid culture and AdenoCre-mediated *Nnmt* knockout *in vitro*. (C) PCR amplification showing *Nnmt* knockout allele using genomic DNA from the indicated organoids. (D) TaqMan qRT-PCR confirming NNMT loss in *Nnmt^-/-^* organoids. (E) Quantification of cell numbers in organoid culture experiments using the indicated cells as shown in Figure 1B. (F) Prostate-specific *Nnmt* knockout in mice. PCR amplification showing *ΔNnmt* fragment in *Nnmt^-/-^* mouse prostate. *Nnmt^flox^* and tail DNA were used as negative controls. (G) Prostate-specific *Nnmt* and *Pten* double knockout in mice. PCR amplification showing ΔNnmt and ΔPten fragments in *Nnmt^-/-^Pten^-/-^* mouse prostate. *Pten^-/-^* and tail DNA were used as negative controls. (H) Pathological categorization of the identified lesions in knockout mice. Representative H&E images of lesions. (I-J) *Pten^-/-^Nnmt^-/-^* mice develop larger prostates at 6 months. (I) Gross image of representative prostate and seminal vesicle (J) Quantification *of* genitourinary weights at 6 and 12 months. (K) Representative IHC images of Smooth muscle Actin (SMA), AR and Ki67 staining showing infiltrating carcinoma developed in *Pten^-/-^Nnmt^-/-^* prostate at 12 months. (L) Quantification of Ki67 staining. Statistical significance was assessed using unpaired two-sided Student’s t-test (*P < 0.05, **P < 0.01, ***P < 0.001 n.s. non-significant).

**Figure S2.**
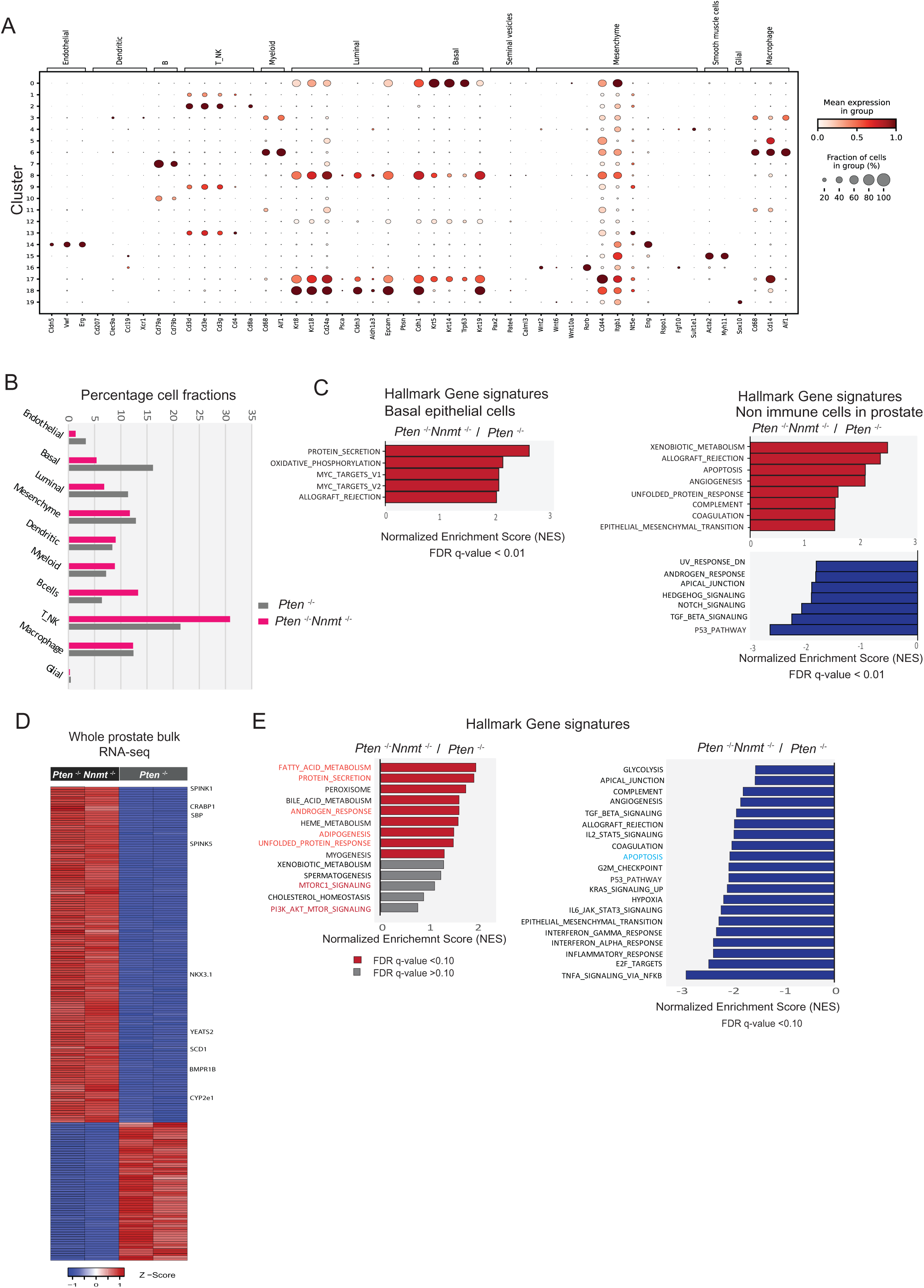
NNMT loss is associated with transcriptional changes. (A) Bubble plot of unique marker genes from different cell types identified through scRNA-seq in *Pten^-/-^Nnmt^-/-^*and *Pten^-/-^* mouse prostate. (B) Distribution fraction of different cell types in *Pten^-/-^Nnmt^-/-^* and *Pten^-/-^* prostate. (C) List of significant Hallmark gene signatures in basal epithelial and non-immune cells from *Pten^-/-^Nnmt^-/-^* versus *Pten^-/-^* prostate. Note the absence of mTOR and AR gene signatures in these cell population unlike luminal epithelium shown in Figure 2. (D) Heatmap of differentially expressed genes derived from RNA-seq of *Pten*^-/-^*Nnmt*^-/-^ and *Pten*^-/-^ whole prostate tissue from mice fed with methionine-restricted diet. (E) Enriched and depleted Hallmark gene signatures in *Pten^-/-^Nnmt^-/-^*prostates from mice fed with methionine-restricted diet.

**Figure S3.**
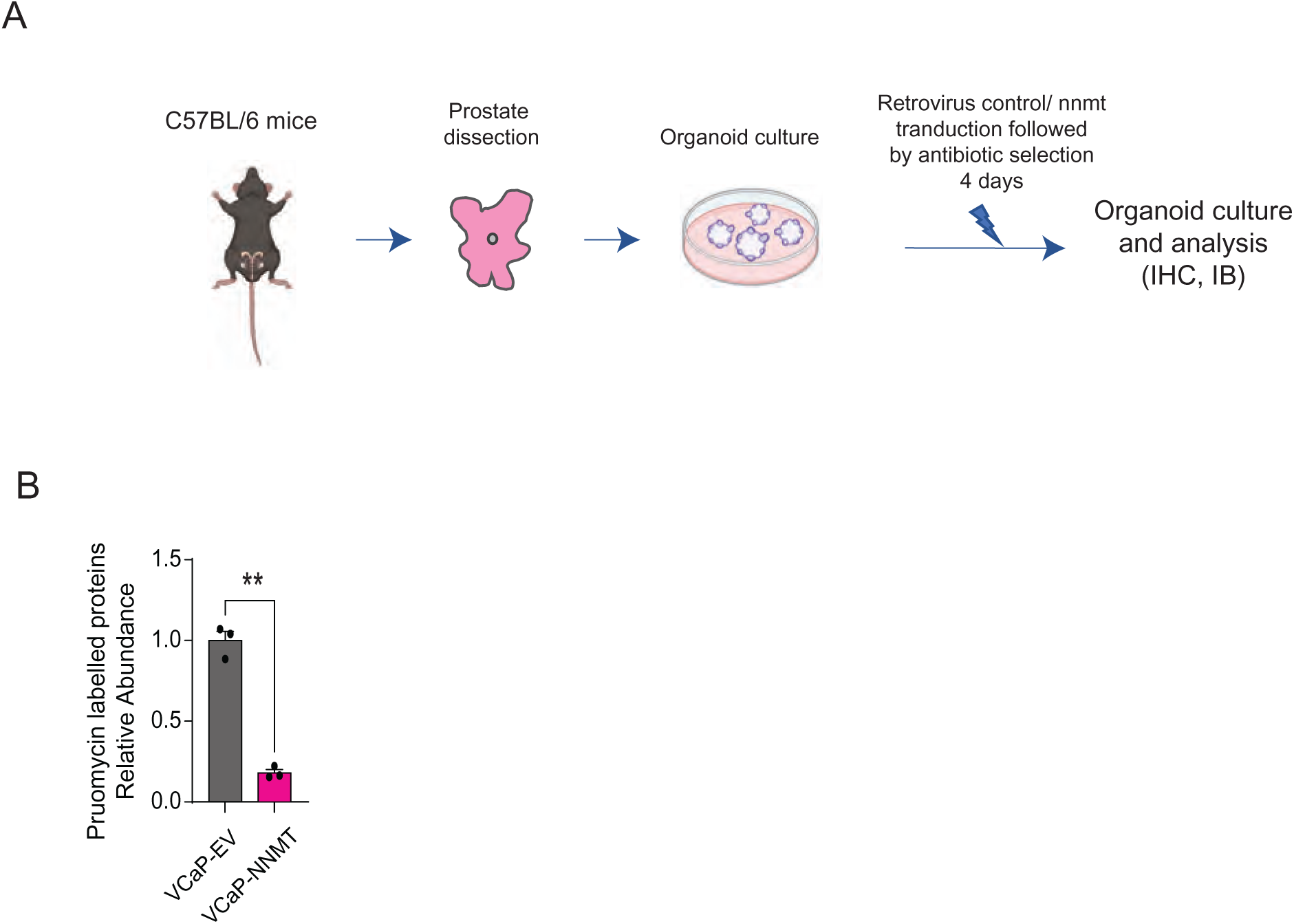
NNMT modulates mTORC1 signaling and protein translation. (A) Schematic of prostate organoid culture and retrovirus based NNMT overexpression. (B) Quantification of the immunoblot signal from Figure 3H. Statistical significance was assessed using a paired two-sided Student’s t-test (**P < 0.01).

**Figure S4.**
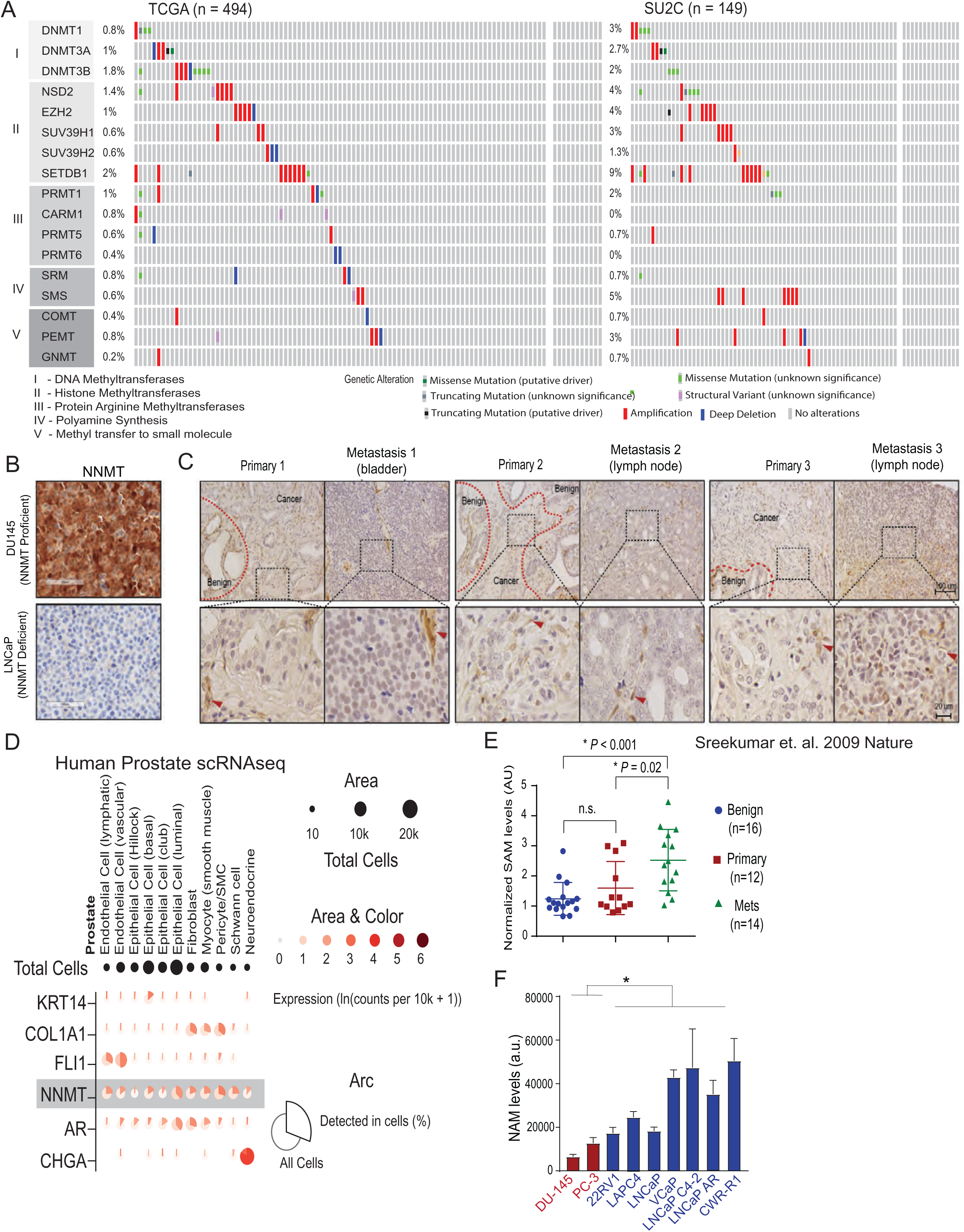
Recurrent genomic deletion and silencing of NNMT in human prostate cancer. (A) Oncoprint from CBioPortal showing genetic alterations in major SAM-consuming enzymes in primary (TCGA) and metastatic (SU2C) prostate cancer datasets. (B) NNMT antibody validation DU145 and LNCaP cells. (C) NNMT expression is lost in human PCa. IHC images showing NNMT expression in matched primary and metastatic human PCa tissue samples. The lower panels are magnified selected regions and red arrows indicate NNMT-positive non-cancerous cells. (D) NNMT expression in normal human prostate. The bubble plot illustrating the expression levels of NNMT alongside other cell type-specific genes across various cell types within the normal prostate, as derived from GTEx Portal. (E) Elevated SAM levels in prostate cancer metastases. LC-MS quantification of SAM levels in benign, localized PCa, and metastatic castration-resistant prostate cancer (CRPC) tissues. The plot was generated from data within the published metabolome datasets (PMID: 19212411). (F) LC-HRMS-based quantification of nicotinamide (NAM) levels in PCa cell lines.

**Figure S5.**
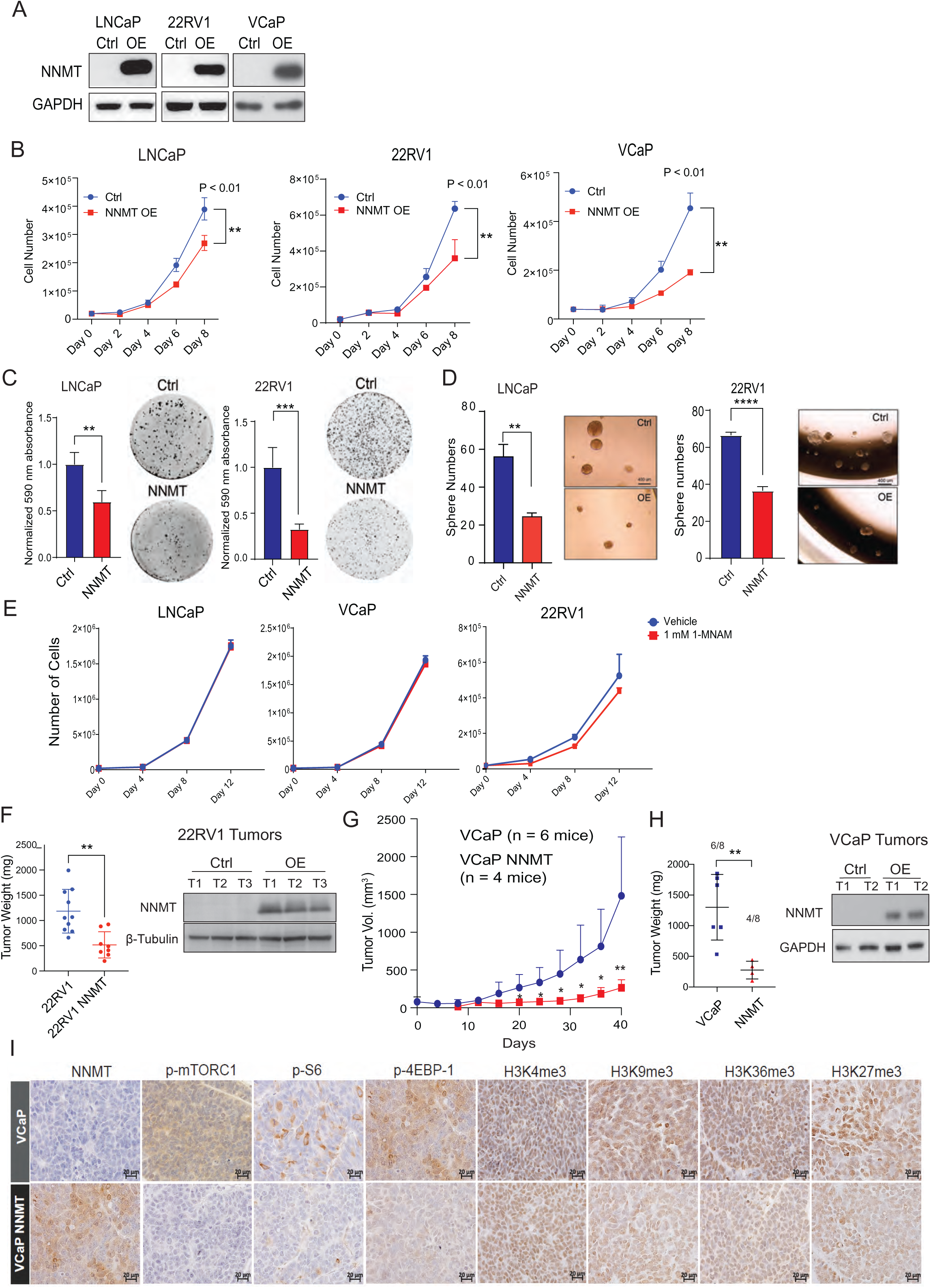
NNMT overexpression suppresses prostate cancer growth *in vitro* and *in vivo*. (A) NNMT overexpression models in PCa cells. Immunoblots confirm NNMT expression in engineered cells. (B-D) NNMT overexpression suppress PCa growth *in vitro*. (B) Cell proliferation rates over time. (C-D) Quantifications and representative images of colony (C) and sphere (D) formation assays. Cells were grown in regular media for three assays. (E) 1-MNAM has no effect on prostate cancer cell proliferation. Growth rates of LNCaP, VCaP, and 22RV1 cells cultured with 1mM 1-MNAM or vehicle over time. (F-H) NNMT overexpression suppresses AR-positive PCa xenograft growth in mice. (F) Endpoint 22RV1 tumor weight from main Figure 4F. Immunoblots confirming NNMT overexpression in xenograft lysates. (G,H) As in F with VCaP and VCaP NNMT overexpression model. (I) Reduced mTOR signaling and histone methylation in NNMT-overexpressing VCaP xenografts. Representative IHC images of the phospho-proteins in mTORC1 pathway and indicated histone methylation marks in control and NNMT-overexpressing VCaP tumor xenografts. Statistical significance was assessed using an unpaired two-sided Student’s t-test (*P < 0.05, **P < 0.01, ***P < 0.001 ****P < 0.0001).

**Figure S6.**
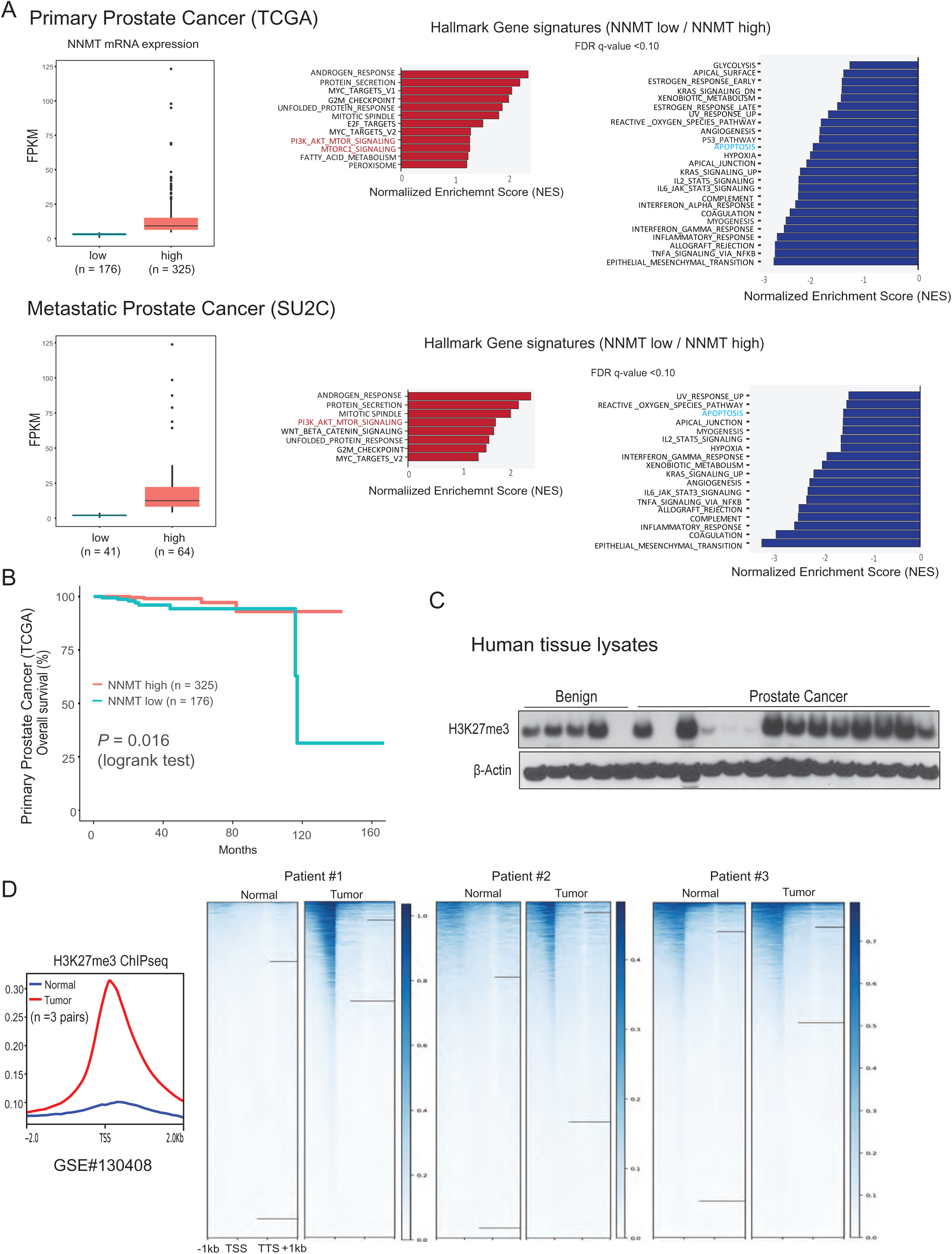
Altered gene expression pathways and histone methylation in human PCa. (A) Top-ranked significantly enriched and depleted Hallmark gene signatures in NNMT-low versus NNMT-high tumors from TCGA and SU2C prostate cancer datasets. Box plots show NNMT expression in low and high cases. (B) Kaplan-Meier survival curves for NNMT low and high cases in TCGA dataset. (C) Immunoblots of histone H3K27me3 marks in human PCa tumor and benign prostate tissue lysates. b-Actin served as loading control. (D) *left,* H3K27me3 ChIP-seq in matched normal and PCa, reanalyzed from GSE1304408. *Right,* heatmap representation across +/- 2Kb TSS and TTS.

**Figure S7.**
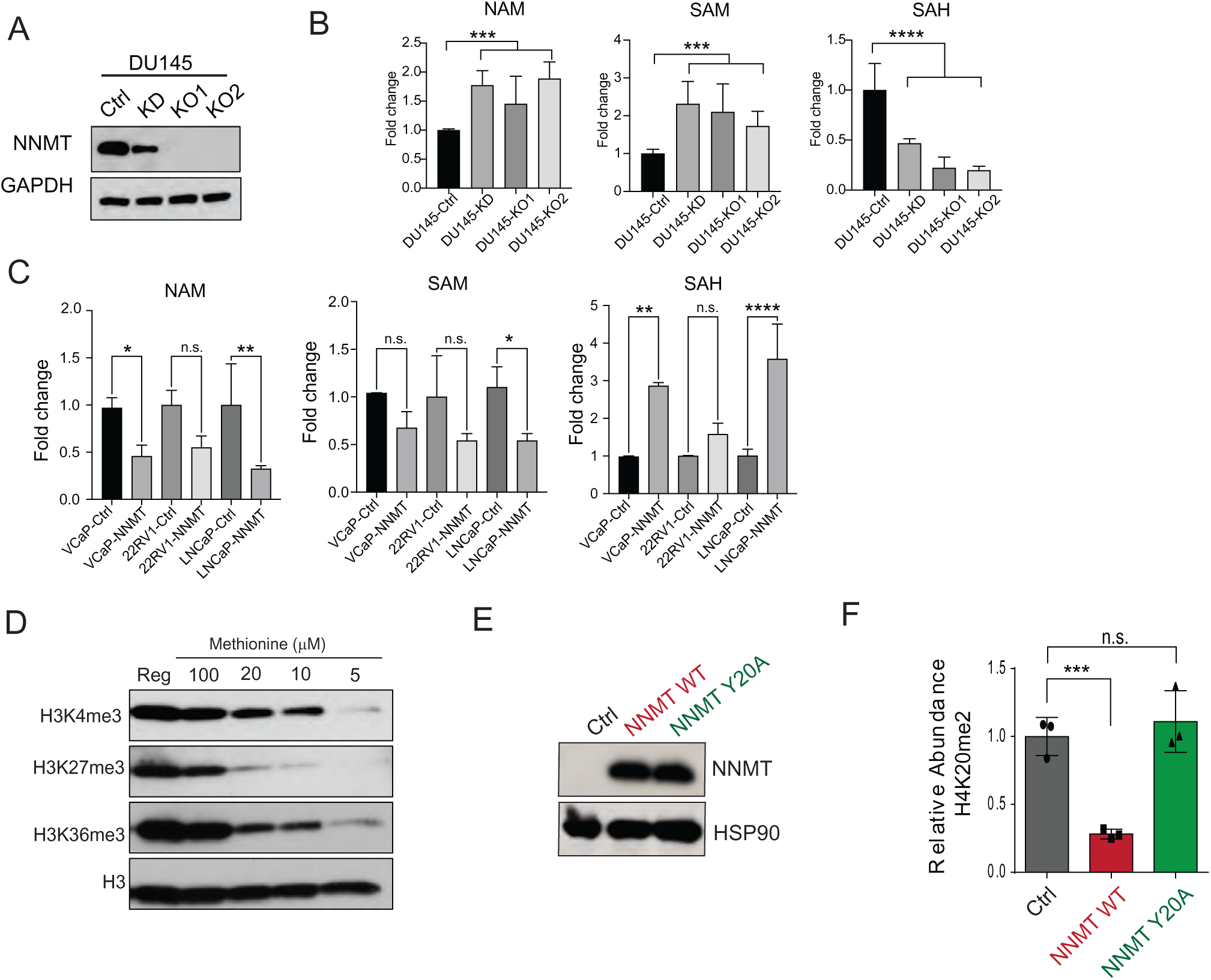
NNMT catalytic activity regulates cellular SAM levels, while MNAM does not influence prostate cancer cell growth. (A) Establishment of NNMT knockdown and knockout models. Immunoblots confirm NNMT depletion. (B-C) NNMT depletes cellular SAM levels. LC-MS analysis shows reciprocal changes in NAM, SAM, and SAH levels in NNMT KO and overexpressing prostate cancer cells. (D) DU145 cells grown in the indicated methionine concentration for 12-days. Histone extracts were probed with the indicated antibody. (E) Immunoblots of lentivirus-based wildtype NNMT or the NNMT Y20A catalytic-dead mutant expression in LNCaP cells. (F) Quantification of H4K20me2 immunoblot shown in Figure 5H. Statistical significance was assessed using an unpaired two-sided Student’s t-test (*P < 0.05, **P < 0.01, ***P < 0.001 ****P < 0.0001, n.s. non-significant).

**Figure S8.**
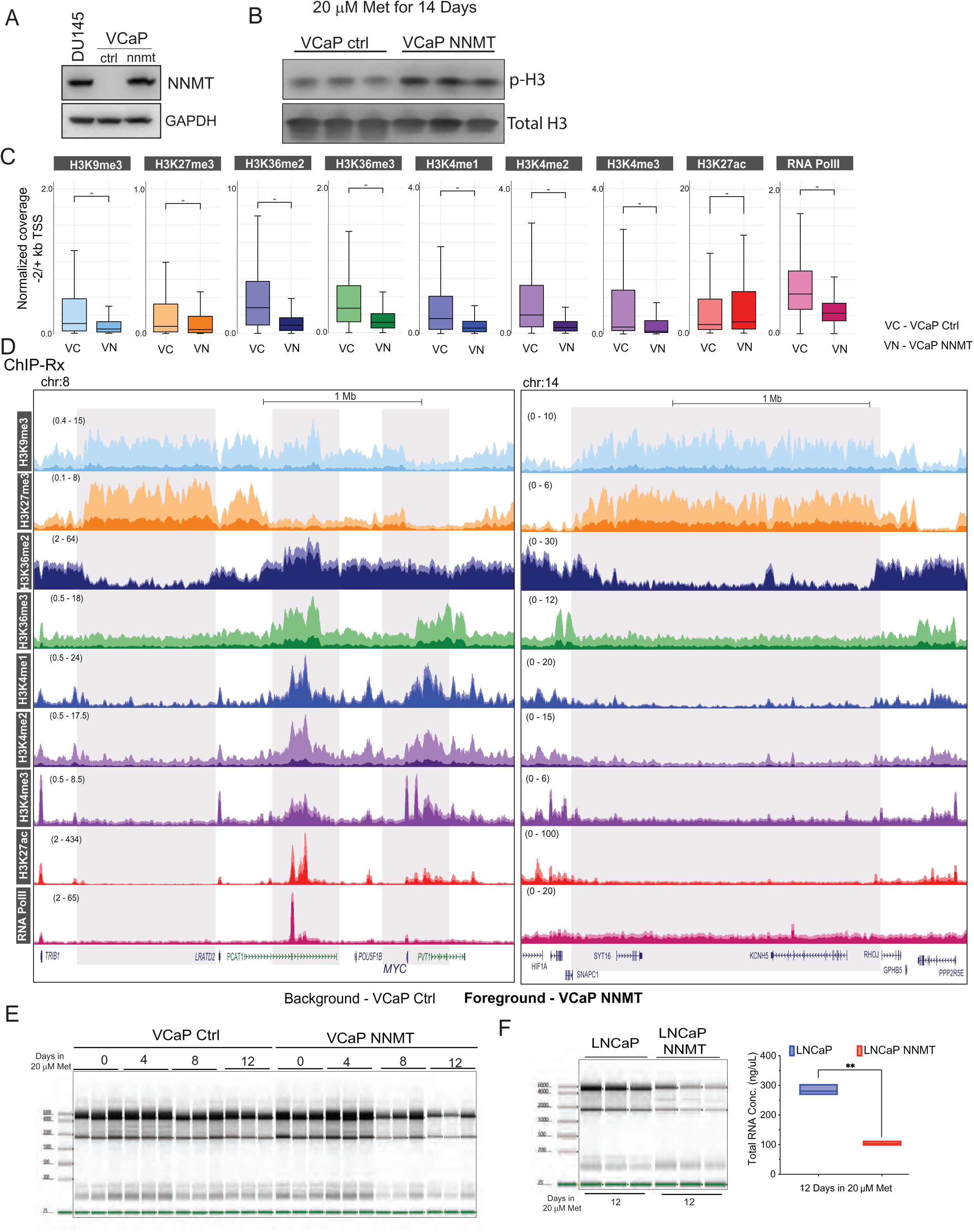
NNMT depletes histone methylation genome-wide and suppresses transcription in prostate cancer cells. (A) Immunoblots of NNMT in DU-145, VCaP, and VCaP NNMT overexpressing cells. (B) Immunoblot of phospho-H3, a surrogate for mitotic cells, in VCaP control and NNMT-overexpressing cells, grown in physiological concentration of methionine for 14-days. (C-D) NNMT expression decreases histone methylation. (C) Histone methylation and acetylation marks and RNA PolII at TSS (+/- 2kb) (D) Genome browser view of transcriptionally active (*MYC* locus) and silenced (*KCNH5* locus) regions. The background lighter shade signal is from VCaP control, and the darker shade signal in the foreground is from VCaP NNMT overexpressing cells. The gray box indicates mutually exclusive hetero- and euchromatin regions. Data normalized with drosophila chromatin spike-in controls (ChIP-Rx). (E-F) NNMT expression suppresses the transcription rate in PCa cells. (E) TapeStation gel images for RNA from VCaP control and NNMT overexpressing VCaP cells grown under physiological methionine conditions – as in Fig.5I. (F) TapeStation gel image with total RNA and its concentrations from NNMT-overexpressing LNCaP cells. Statistical significance was assessed using an unpaired two-sided Student’s t-test, **P < 0.01.

**Figure S9:**
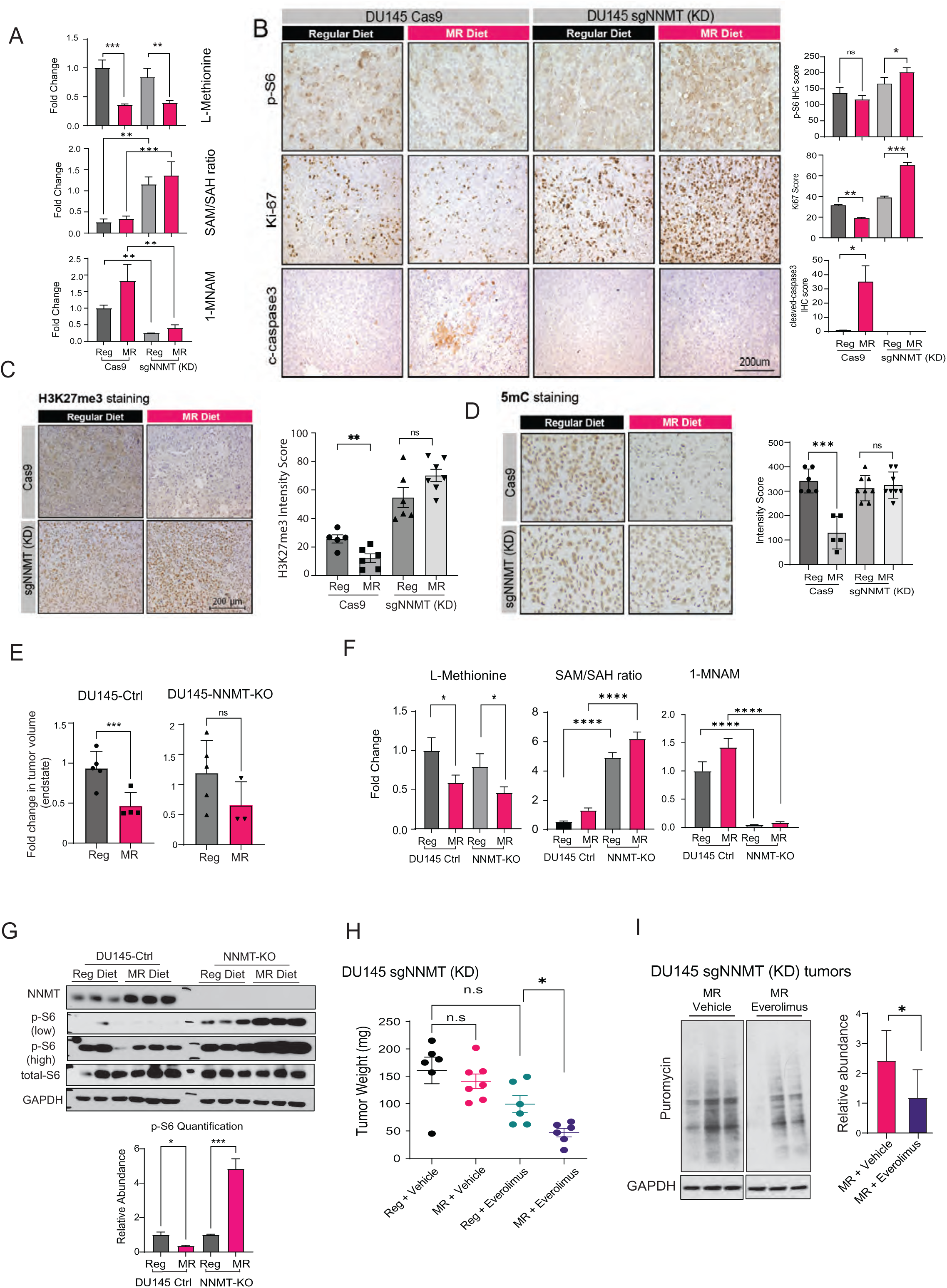
NNMT deficiency creates therapeutic vulnerability to mTOR inhibitor when combined with methionine restricted diet. (A) MR-diet reduces methionine levels in xenograft tumors. LC-MS quantification of methionine, SAM/SAH ratio, and 1-MNAM in DU145 cas9 control and DU145 NNMT KD tumor lysates. (B) MR-diet induces p-S6 and mitotic activity in the NNMT KD tumors. Representative IHC images and quantifications. (C-D) Compared to NNMT-proficient controls NNMT KD tumors show no reduction in H3K27me3 and 5-methylated Cytosine (5mC) levels under MR-diet. Representative IHC images and quantifications. (E-G) NNMT-knockout DU145 tumors are refractory to MR-diet through activation of mTOR signaling. (E) Endpoint tumor volume under MR (n = 4) or regular diet (n=5). (F) LC-MS quantification of SAM metabolites. (G) Immunoblots and quantifications of p-S6. (H) MR-diet synergizes with Everolimus (mTOR inhibitor) to suppress DU145 NNMT-KD tumors. Endpoint tumor weights in DU145 NNMT knockdown xenografts as shown in Figure 6F. (I) *In vivo* SUnSET assay shows reduced protein synthesis by MR and Everolimus treatment in NNMT-KD DU145 xenografts. *Left,* immunoblots of puromycin using the indicated xenograft lysates, each lane represents an independent tumor. *Right,* Quantifications. Statistical significance was assessed using an unpaired two-sided Student’s t-test (*P < 0.05, **P < 0.01, ***P < 0.001 ***P < 0.0001, n.s. non-significant).

**Figure S10:**
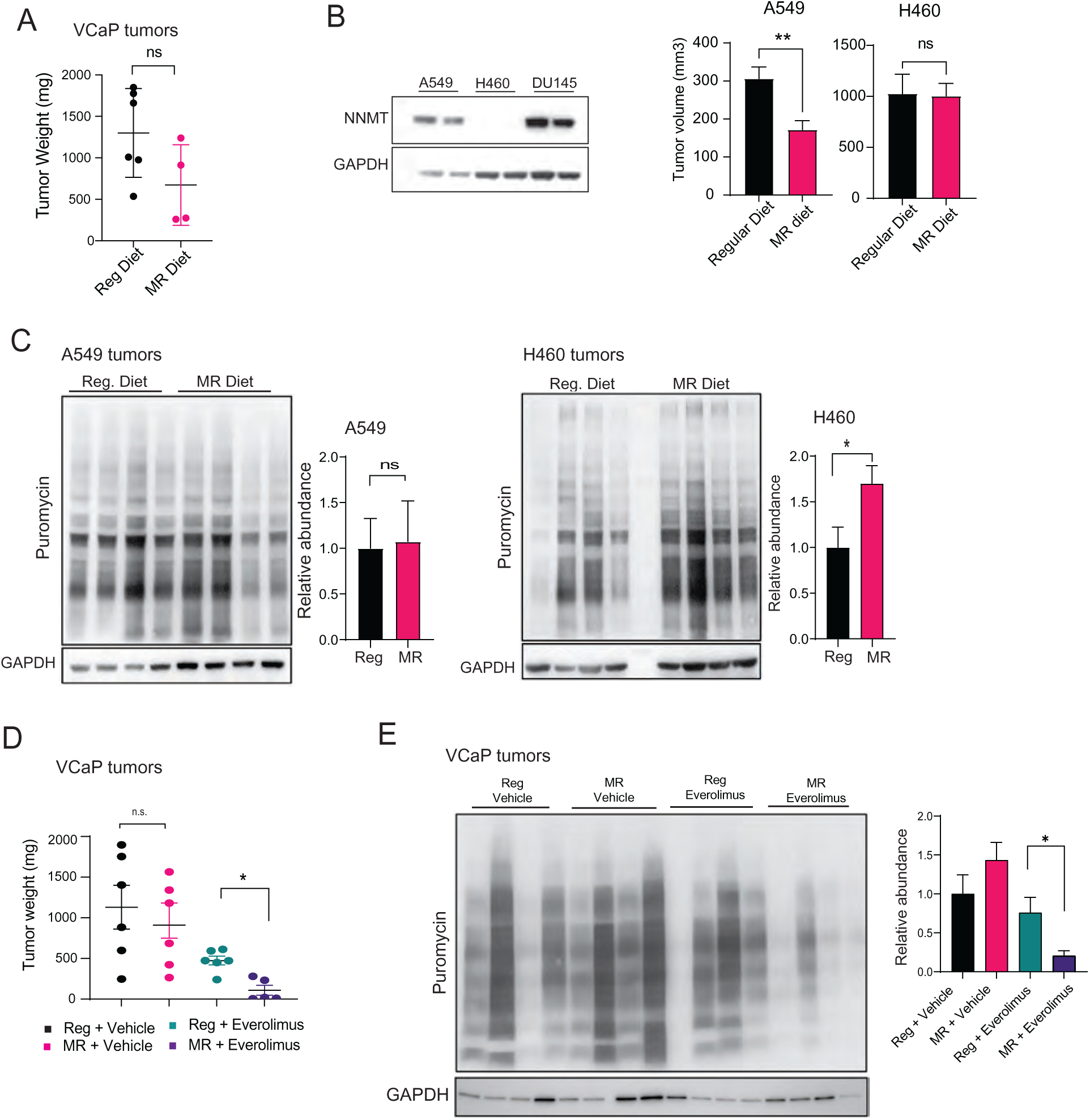
NNMT deficiency creates therapeutic vulnerability to mTOR inhibitor when combined with methionine restricted diet. (A) NNMT-deficient VCaP xenografts are refractory to MR-diet. Endpoint tumor weight of a VCaP xenograft assay in mice fed with regular (n = 6) or MR-diet (n = 4). (B-C) NNMT-deficient human lung xenografts are refractory to MR-diet (B) *Left,* immunoblots of NNMT in human lung cancer cell lines: A549 and H460 and PCa cell line DU145. *Right,* endpoint tumor volume in NNMT-proficient A549 and NNMT-deficient H460 xenograft assays in mice fed with regular or MR-diet. (C) Immunoblots of puromycin using the indicated xenograft lysates, each lane represents an independent tumor. *Right*, Quantifications (D-E) MR-diet combined with Everolimus suppresses VCaP xenograft growth. Endpoint VCaP xenograft tumor weights and immunoblots of puromycin using the indicated xenograft lysates, each lane represents an independent tumor. *Right*, Quantifications Statistical significance was assessed using an unpaired two-sided Student’s t-test (*P < 0.05, **P < 0.01, n.s. non-significant).

## Key Resources Table

Reagent or resource, source, identifier (cat #)

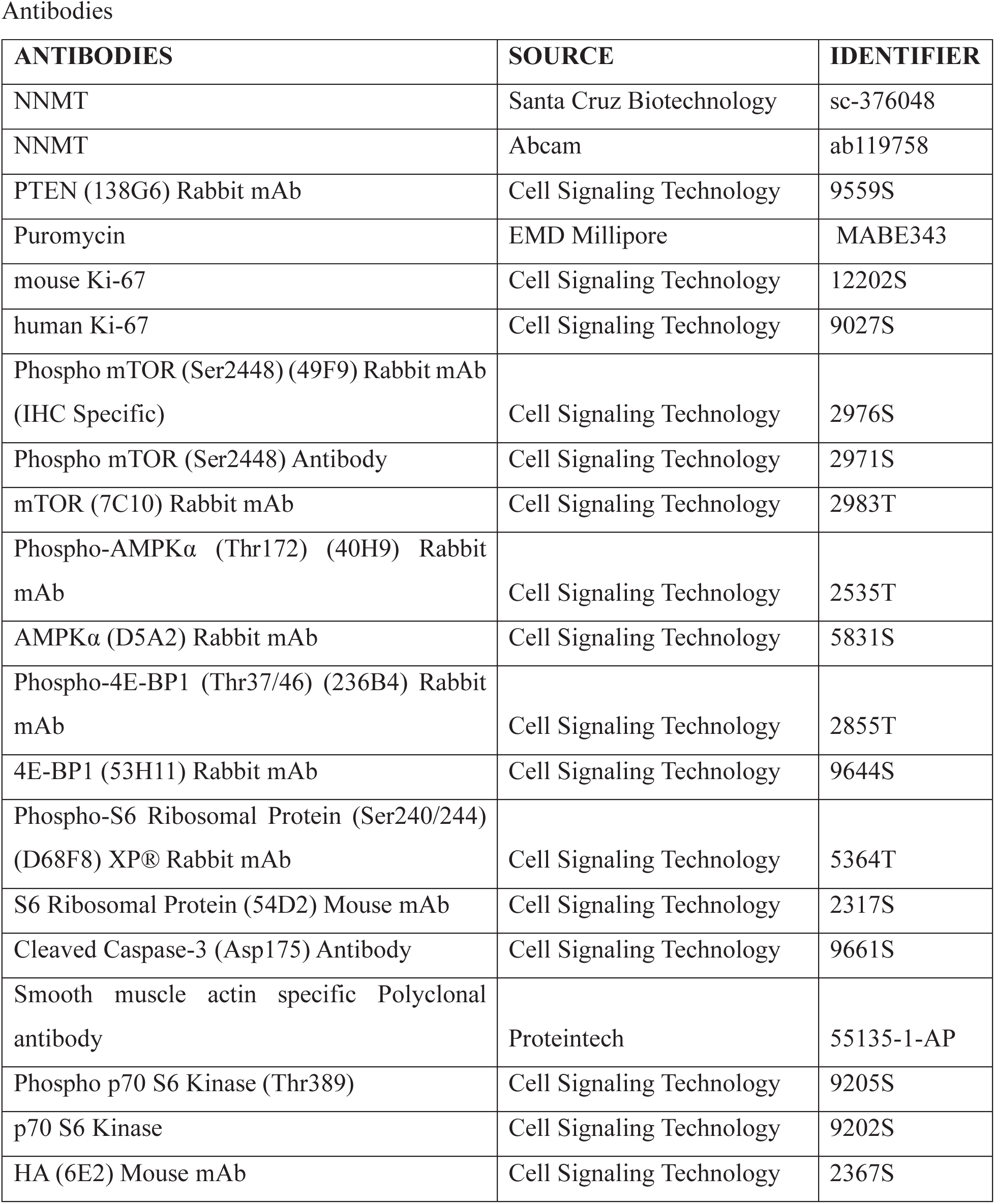
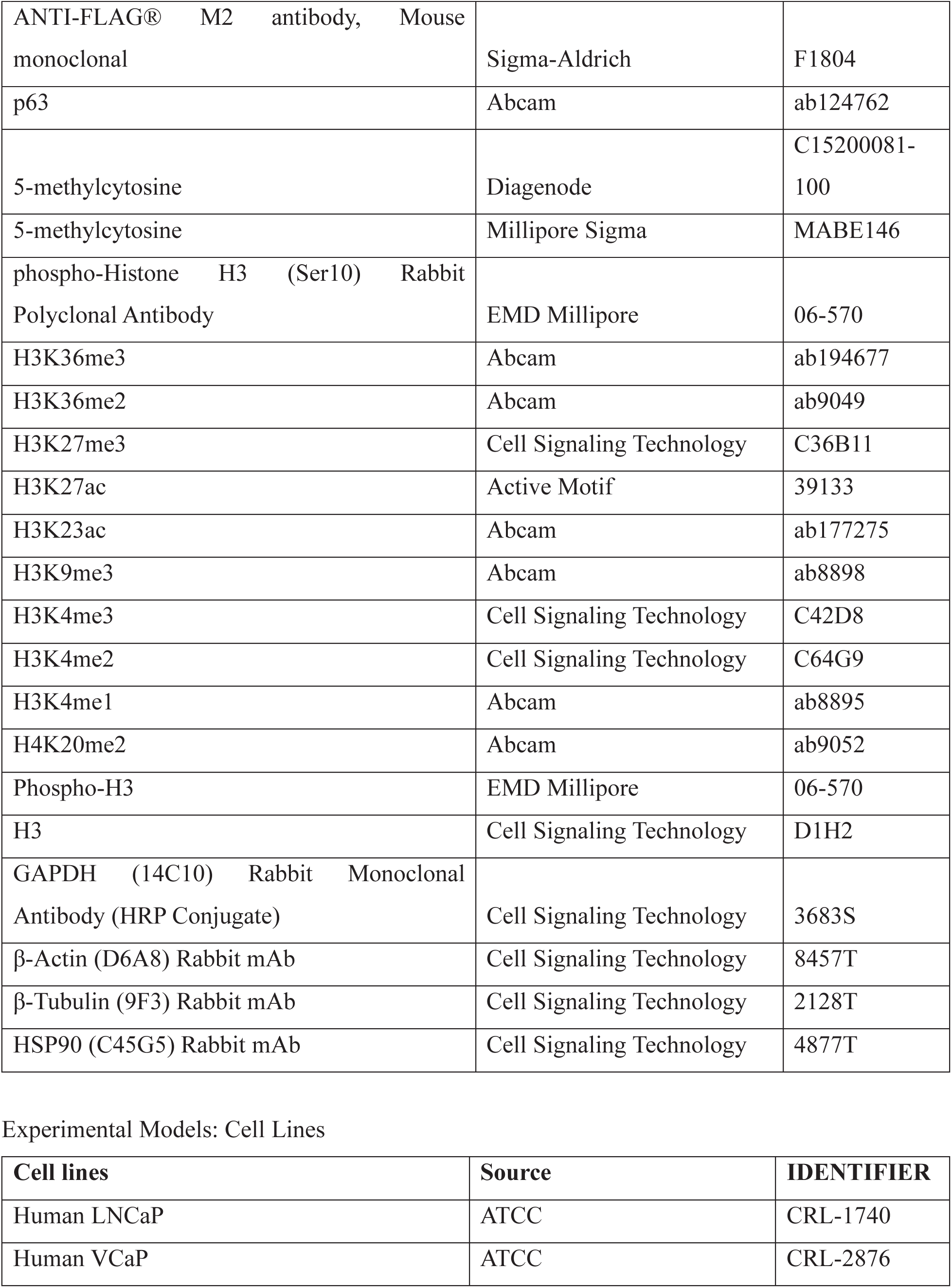
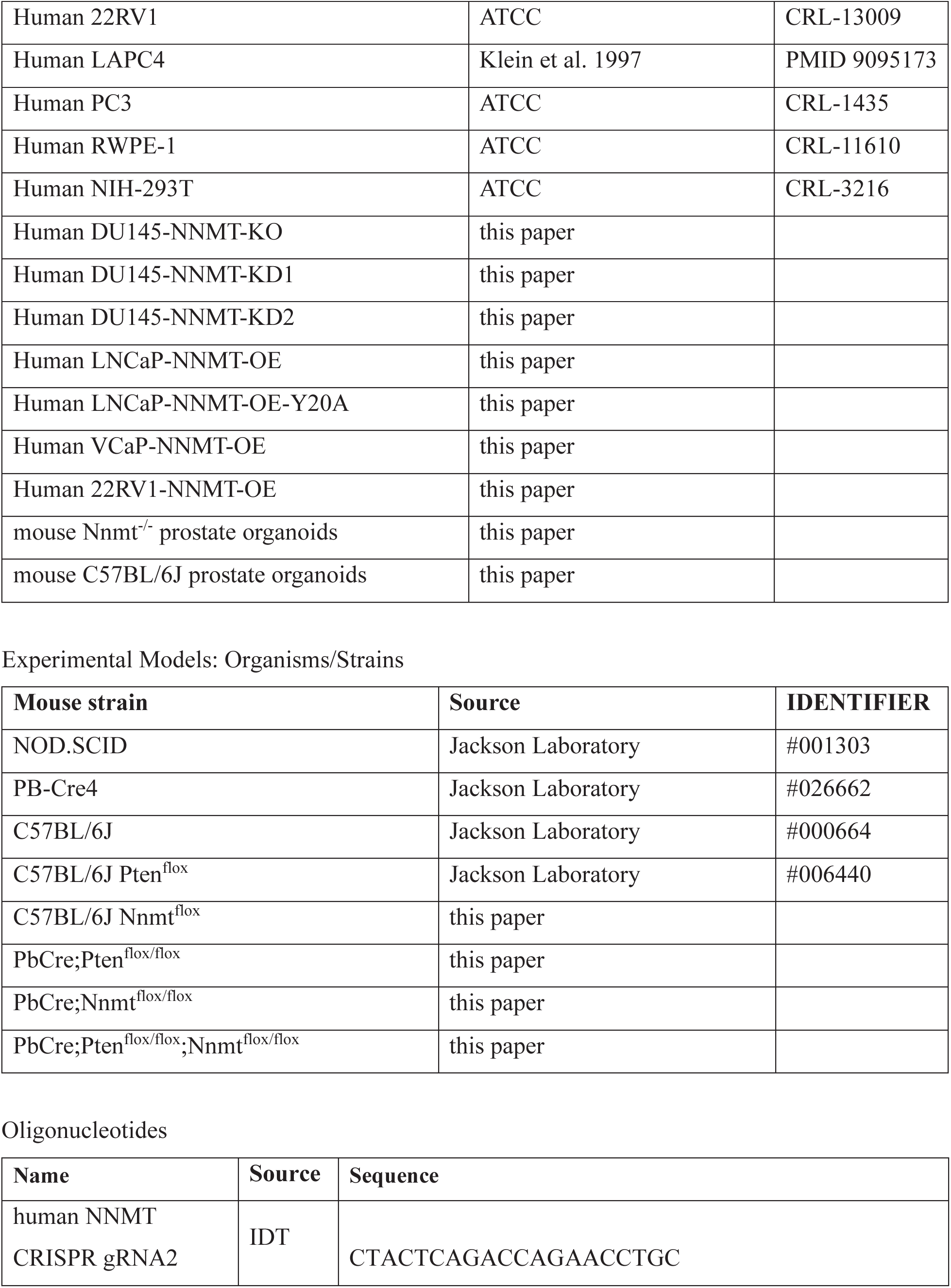
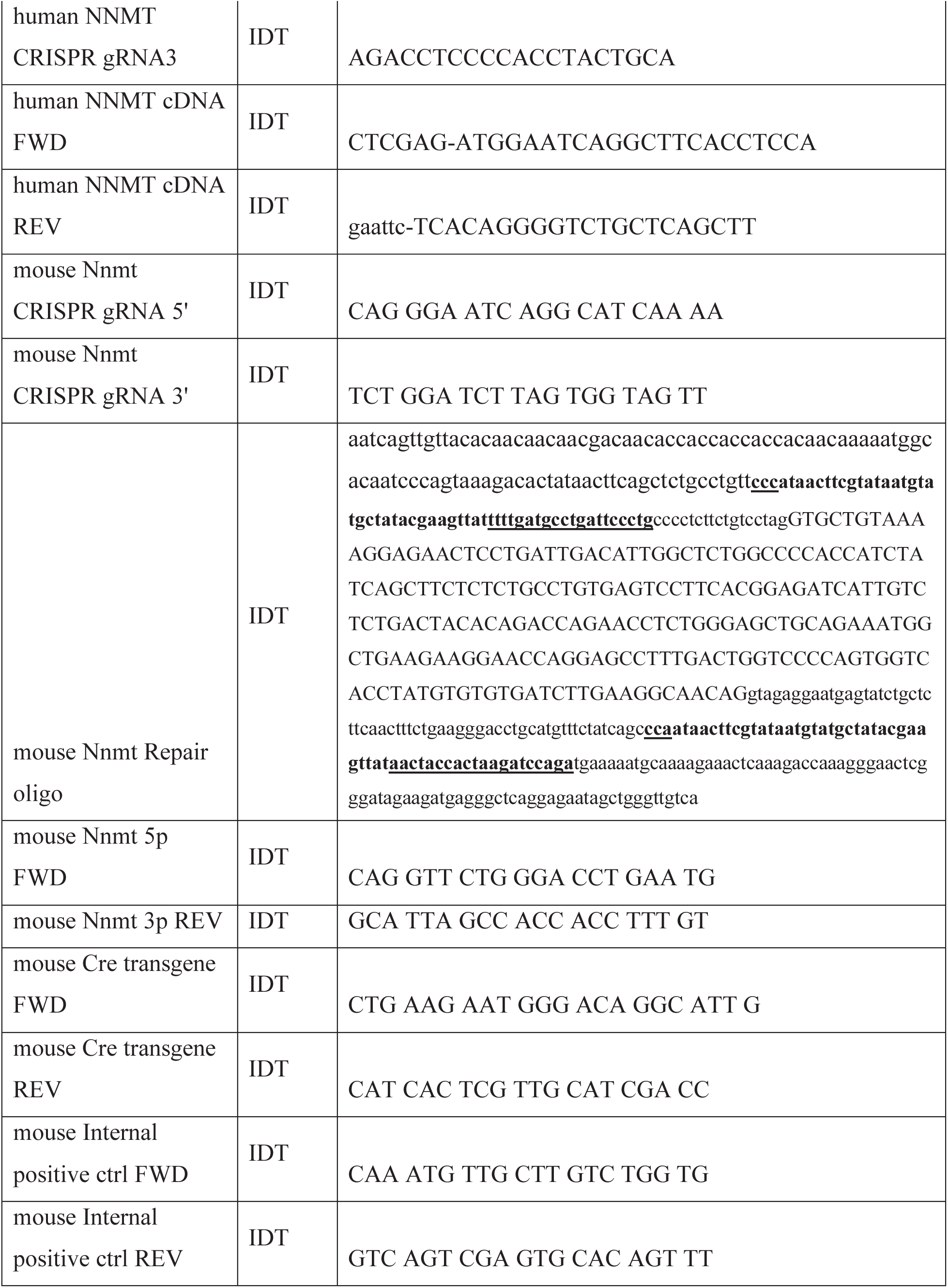
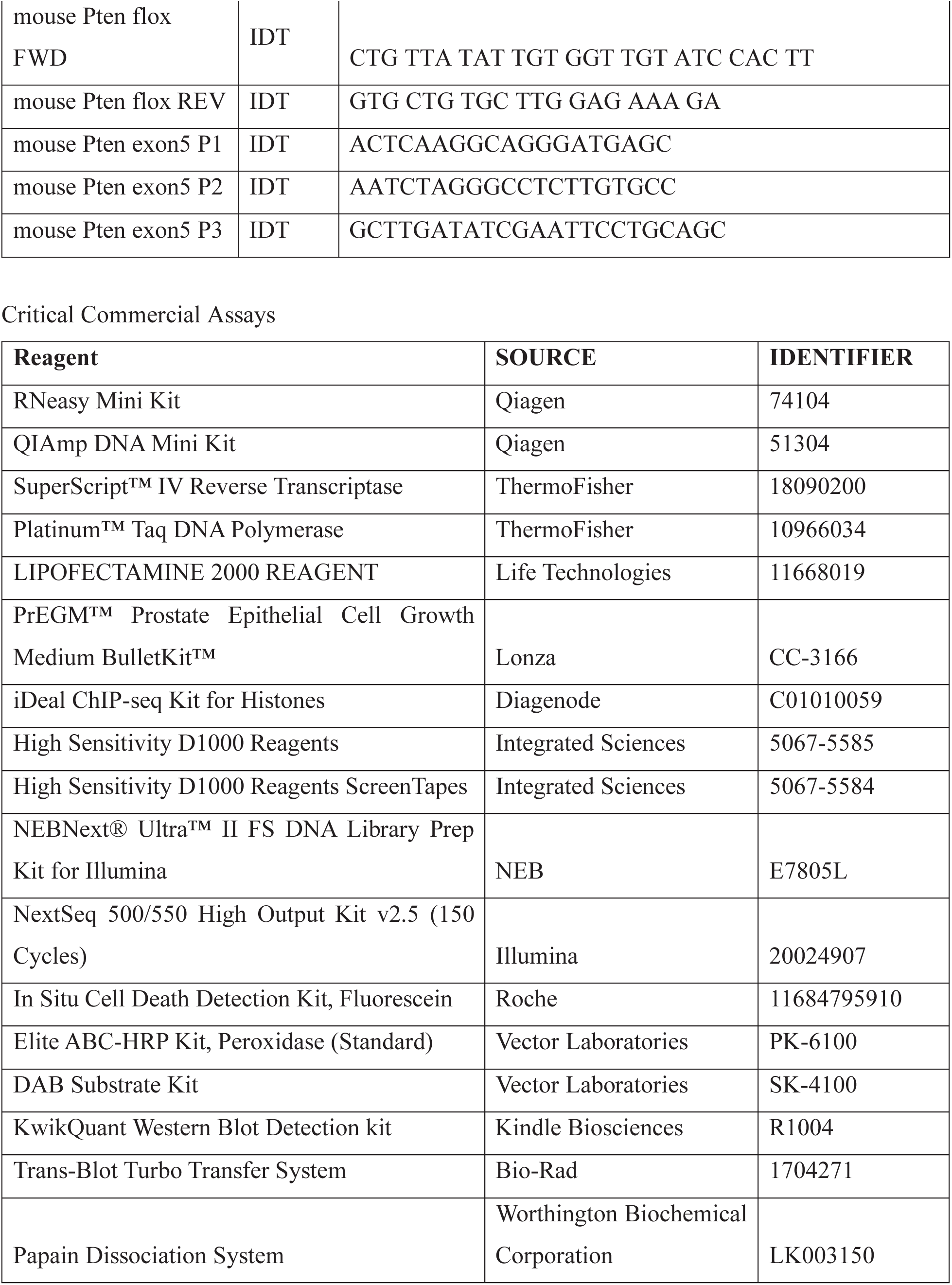
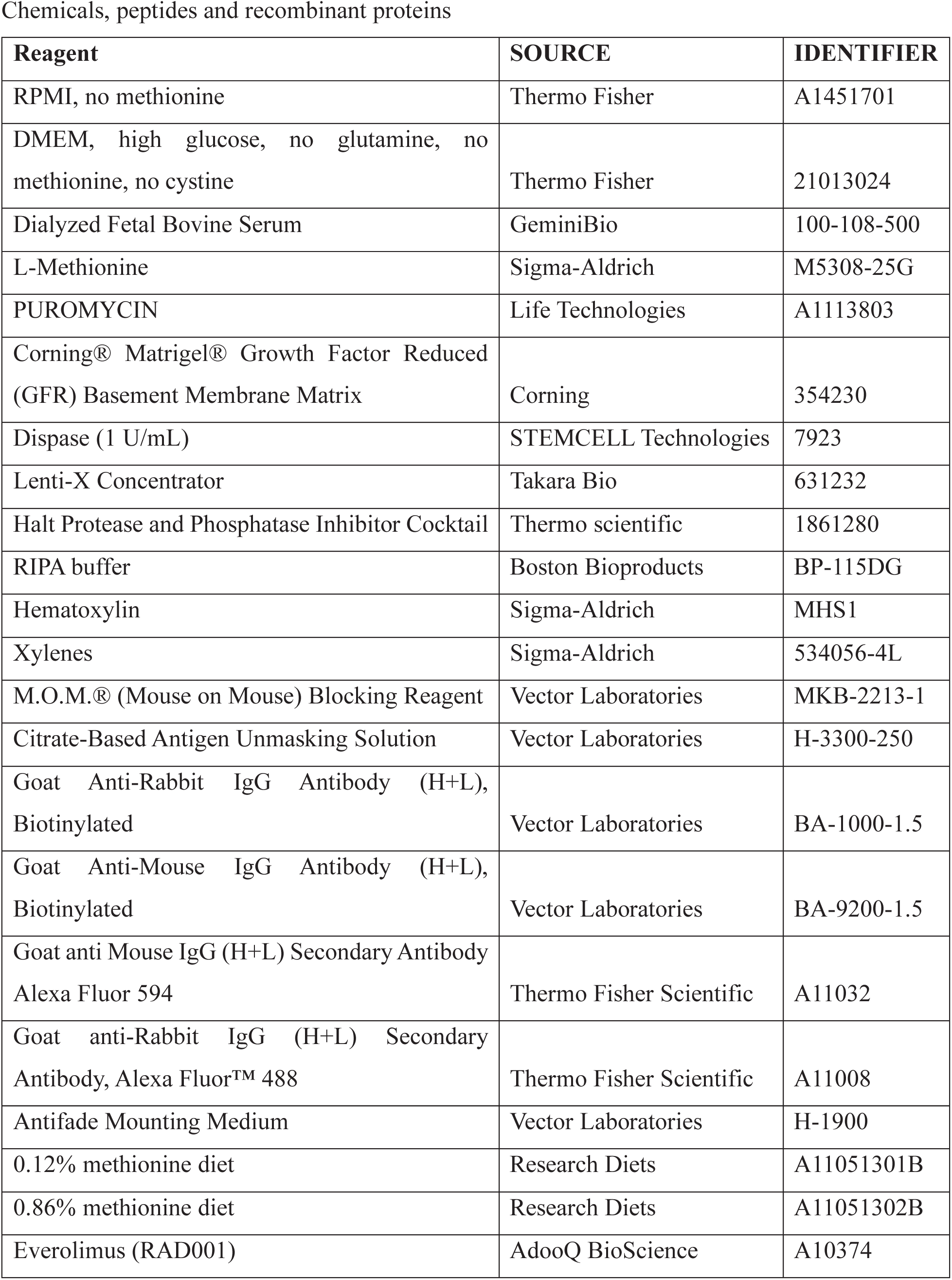
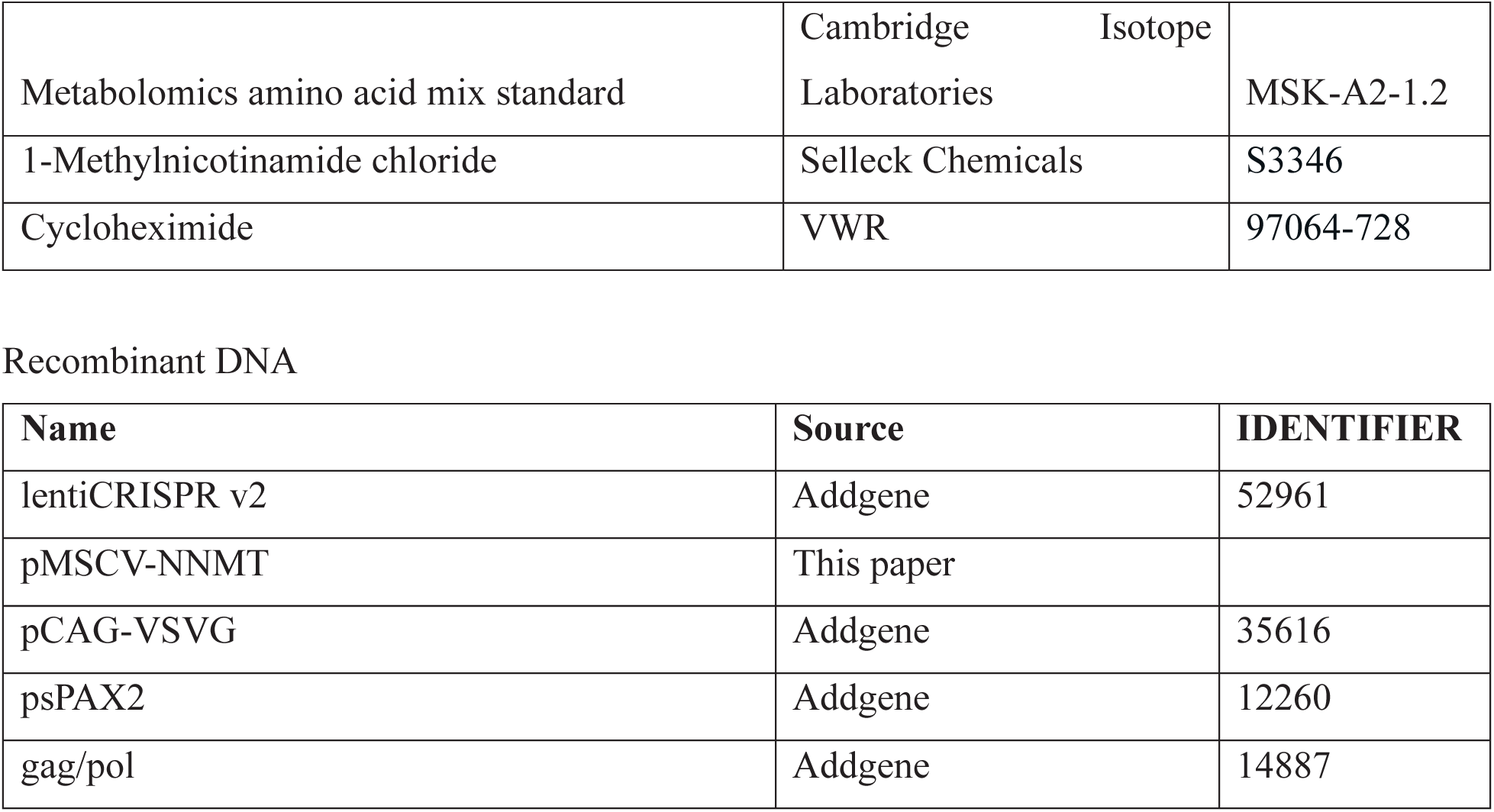

Software and Algorithms

PRISM 10, GraphPad, Version 10.2.2

## Resource and Materials Availability

-Lead Contact: Irfan A. Asangani

## -Data and Code Availability

All sequencing data generated in this study have been deposited in the Gene Expression Omnibus under accession **GSE285539** (n=32, **access token: slajkocijderxkb**). The current submission includes 22 ChIP-seq samples, 8 bulk RNA-seq samples, and 2 single-cell RNA-seq datasets, with additional metadata provided as required.

## Experimental model and subject details

### Human prostate cancer tissue microarray

In the selection of prostate tissues for a tissue microarray (TMA) encompassing benign, primary prostate cancer, and metastatic prostate cancer samples, archived formalin-fixed paraffin-embedded (FFPE) tissue blocks were initially assessed by skilled pathologists. Representative regions from benign prostatic tissue, primary prostate cancer lesions, and metastatic deposits (from distant sites like lymph nodes or bone) were identified through histological examination. Tissue cores, typically 0.6 to 2 mm in diameter, were precisely extracted from these diverse regions of interest within each tissue block using a tissue microarray instrument. These extracted cores were meticulously arranged in a pre-designed layout within a new paraffin block, thereby creating the tissue microarray. Subsequently, the TMA block was sectioned into thin slices of 3-5 micrometers using a microtome, and these sections were mounted onto glass slides for further immunohistochemical analysis. Glandular NNMT staining was scored by two pathologists, independently.

### Genetically engineered mouse models

Animal studies were performed in accordance with protocols approved by the Institutional Animal Care and Use Committee at the University of Pennsylvania and in compliance with all regulatory standards. To generate the Nnmt conditional allele, CRISPR gRNAs: 5’ CAG GGA ATC AGG CAT CAA AA and 3’ TCT GGA TCT TAG TGG TAG TT and a repair oligo were designed to knock in two loxP sites flanking the mouse *Nnmt* exon 2. The founder *Nnmt ^flox^* mice were backcrossed on C57BL/6 mice for a minimal two generations to clean the background. *Nnmt ^flox/flox^* mice were crossed with *Pten^flox/flox^* (B6.129S4-Ptentm1Hwu/J Strain #:006440) mice to generate the *Nnmt ^flox/flox^*;*Pten^flox/flox^* mice.

The male Pbsn-Cre mice (Tg(Pbsn-cre)4Prb/J, Jackson laboratory: Strain #:026662) were crossed with female *Nnmt ^flox/flox^*, *Pten^flox/flox^* and *Nnmt ^flox/flox^*;*Pten^flox/flox^* mice to generate the prostate specific knockout of the flox alleles. Genotype of the mice were confirmed using the following primers: Nnmt 5p Forward: CAG GTT CTG GGA CCT GAA TG, Nnmt 3p Reverse: GCA TTA GCC ACC ACC TTT GT. Pten flox Forward: CTG TTA TAT TGT GGT TGT ATC CAC TT; Pten flox Reverse: GTG CTG TGC TTG GAG AAA GA. Cre genotyping primers: Transgene Forward: CTG AAG AAT GGG ACA GGC ATT G, Transgene Reverse: CAT CAC TCG TTG CAT CGA CC, Internal Positive Control Forward: CAA ATG TTG CTT GTC TGG TG, Internal Positive Control Reverse: GTC AGT CGA GTG CAC AGT TT.

### Cell lines and cell culture

Prostate parental cell lines used in this study were obtained from ATCC. LNCaP, 22RV1, LAPC4, DU145, PC3 prostate cancer cell lines were grown in RPMI-1640 (Gibco, 11875093), and the VCaP cells was grown in DMEM with Glutamax (Gibco, 21013024). The medium was supplemented with 10% of FBS (HYC, SH30910.03) and 1% of penicillin–streptomycin solution (Invitrogen, 15140122). The immortalized benign prostate cell line RWPE-1 was grown in keratinocyte media with supplements (Thermo Fisher Scientific, 17005042). The cell lines tested negative for Mycoplasma contamination and were maintained in a humidified incubator at 37°C and 5% CO2.

For experiments involving methionine restriction, LNCaP and DU145 cells were cultured in methionine deficient RPMI (Thermo Scientific, A1451701) and VCaP cells were cultured in methionine deficient DMEM (Thermo Scientific, 21013024). The medium was supplemented with either 20 μM or 100 μM methionine (Sigma Aldrich, M5308-25G), 10% of dialyzed FBS (GeminiBio, 100-108), and 1% of penicillin–streptomycin solution (Invitrogen, 15140122). The cell lines tested negative for Mycoplasma contamination and were maintained in a humidified incubator at 37°C and 5% CO2.

## Method Details

### NNMT overexpression

DU145 cell RNA was extracted using RNeasy Mini Kit and cDNA was synthesized using SuperScript™ IV Reverse Transcriptase. Full length human NNMT cDNA (795bp) was cloned from DU145 cell cDNA using Platinum™ Taq DNA Polymerase and the following primers: XhoI-FWD: CTCGAG-ATG GAA TCA GGC TTC ACC TCC A EcoRI-REV: GAATTC – TCA CAG GGG TCT GCT CAG CTT. The fragment was then ligated into a retrovirus backbone: pMSCV-puro (Takara, 631461) using restriction enzyme cutting sites XhoI and EcoRI. To generate a catalytically inactive NNMT mutant, a tyrosine-to-alanine substitution at residue 20 (Y20A) was introduced into the NNMT coding sequence by site-directed mutagenesis using the pMSCV-puro-NNMT WT plasmid as a template. The Y20A mutation targets a conserved residue within the NNMT catalytic pocket and has been shown to abolish methyltransferase activity without affecting protein stability. Successful mutagenesis was confirmed by Sanger sequencing across the full NNMT open reading frame. To package for NNMT overexpression retrovirus, 1×10^6^ HEK-293HT cells were seeded in 10cm plates. Next day pMSCV-puro (4 µg) were co-transfected with pVSVg (1 µg) and gag/pol (2 µg) packaging plasmid using 7 µL Lipofectamine 2000. Media collected after 48h and 72h of transfection, centrifuged and passed through 0.45 µm filters to clear of any live cells or debris. Virus was then concentrated by 10X with LentiX concentrator (Takara, 631232) aliquoted and stored at −80°C. LNCaP, VCaP and 22RV1 cells were infected at a MOI of 10 and selected with puromycin at 1µg/ml for three days to obtain stably overexpressed cells. Cells within 5 passages after puromycin selection were used for all the experiments.

### CRISPR-cas9 mediated Nnmt knockout

sgRNAs targeting human NNMT were designed with the following website: https://portals.broadinstitute.org/gpp/public/analysis-tools/sgrna-design. For the enhancer region knockout, two pairs of sgRNAs (gRNA2: CTA CTC AGA CCA GAA CCT GC gRNA3: AGA CCT CCC CAC CTA CTG CA) were inserted into lentiCRISPR v2 (Plasmid #52961) backbone. Lentivirus was packaged using 2nd generation lentiviral packaging systems using the following protocol. 1×10^6^ HEK-293HT cells were seeded in 10cm plates. Next day lentiCRISPR (4 µg) were co-transfected with pVSVg (1 µg) and pSPAX2 (2 µg) using 7 µL Lipofectamine 2000. Media collected after 48h and 72h of transfection, centrifuged and passed through 0.45 µm filters to clear of any live cells or debris. Virus was then concentrated by 10X with LentiX concentrator (Takara, 631232) aliquoted and stored at −80°C. DU145 cells were infected at a MOI of 10. Cells were selected with puromycin at 1µg/ml for three days to obtain stably knock-out/down cells. Cells within 5 passages after puromycin selection were used for all the experiments.

### Cell proliferation growth curve

Experiments were performed using 20,000 of each indicated control and NNMT-knockdown or - overexpressing cells were seeded in 24-well plates on day 0, triplicate wells of each group were trypsinized and the cell numbers were counted using Countess II Automated Cell Counter on each indicated timepoint. The regular cultured experiments were performed for day 0, 2, 4 and 6. The hypoxic experiment was performed in 3% O2 for day 0, 2 and 4. For 1-MNAM growth curve, 20,000 of each indicated cell were treated with 1 mM 1-MNAM (Selleck Chemicals, S3346).

### Colony and sphere formation assay

For colony formation assay, 5,000 cells were plated in one well of the 6-well plates, in two weeks cells were fixed and stained with 0.5% of crystal violet, quantification was done by de-staining of crystal violet and measure absorbance at 560nm. For sphere formation assay, 500 cells were suspended in 100µL 50% of Matrigel of full RPMI culture media and spread on the edge of wells in a 24-well plate, we then fill the wells with 1 mL of full RPMI culture media. In three weeks, spheres were counted, and the sizes of the spheres were measured with ImageJ. Triplicates were carried in all the above-described experiments; student t-test was used for statistical analysis.

### Immunoblot analysis

For immunoblot analyses, cells were lysed in RIPA buffer (Boston Bioproducts, BP-115DG) supplemented with protease and phosphatase inhibitor (Thermo scientific, 1861280). Lysates were boiled in SDS sample buffer (Invitrogen) and 30-50 µg of protein was separated by SDS-PAGE and loaded onto a PVDF membrane. Membranes were blocked for one hour in blocking buffer (Tris-buffered saline, 0.1% Tween (TBS-T), 5% non-fat dry milk) and incubated overnight at 4°C in primary antibody. Blots were washed with TBS-T and incubated with HRP-conjugated secondary antibody for one hour at room temperature. Blots were washed again with TBS-T and visualized after incubation with chemiluminescent substrate (GE Healthcare) on Kwik Quant Imager (Kindle Biosciences). The antibodies used in the study are provided in the key resources table.

### TMA histology assessment

Histological features were graded using previously described nomenclature and criteria^71^:

The immunohistochemical stain for NNMT was scored semi quantitatively with negative staining scored as 0, positive staining scored on a scale of increasing intensity of staining from 1+ to 4+. The data was normalized by calculating the H score (percentage of cells positive X intensity of staining)^72^.

### Mouse Prostate Culture

Eight-week-old mice were euthanized, and the prostate gland was dissected in DMEM 10% FBS. Prostate was transferred to a new petri dish with fresh dissecting media and minced with a razor blade and then digested with collagenase on a shaker in 37°C for 2 h. Tissue chunks were span down and further digested with Trypsin/0.05% EDTA for 5 min at 37°C. The cell/tissue suspension was passed through a 40 μm nylon mesh filter to get single cell suspension. 10,000 cells in 40 μL PrEGM media were mixed with 60 μL Matrigel and seeded to the rim of a 12-well plate, followed by adding 800 μL warm PrEGM in 30 min. In 10 days, both wild type and *Nnmt* floxed P0 organoids were digested with Dispase and Trypsin into single cell suspension and treated with Adeno-cre for 20 min to induce NNMT knockout in the *Nnmt* floxed cells. 10,000 treated cells in each group were seeded in 12-well plates, images were taken on day 10 and organoid sizes were measured using ImageJ. Genomic DNA and protein lysates were harvested on day 10 for the indicated molecular analysis.

For NNMT overexpression in mouse prostate organoids, 8-week-old mice were euthanized, and the prostate gland was dissected and processed to obtain single-cell suspensions as described previously^73^. The cells were transduced with a retrovirus encoding NNMT cDNA in single-cell suspension for 30 minutes at 37°C. Following transduction, the cells were seeded in Matrigel and cultured in mouse organoid media composed of advanced DMEM/F12 (adDMEM/F12) supplemented with B27 (1x), N-acetylcysteine (1.25 mM), epidermal growth factor (EGF, 50 ng/mL), Noggin (100 ng/mL), R-spondin 1 (500 ng/mL), A83-01 (200 nM), dihydrotestosterone (DHT, 1 nM), and Y-27632 dihydrochloride (10 μM) during establishment and passaging phases. The organoids were then selected using puromycin (1 μg/mL) for 3 days to ensure successful transduction. After selection, the organoids were expanded, and on day 10, they were harvested for immunohistochemistry and Western blot analysis to evaluate NNMT overexpression and mTORC1 signaling.

### Immunohistochemistry (IHC)

Mouse prostate tissues were fixed in 4% of formaldehyde for 48 h, and paraffin embedded through Molecular Pathology and Imaging Core at UPenn. Paraffin embedded sections from the prostate of indicated genotype were deparaffinized with 3 changes of xylene for 5 min each. Slides were then rehydrated in 100% alcohol for 10 min, 95% alcohol twice for 10 min each, 70% alcohol and distilled water for 10 min each. Slides were subjected to citrate based (pH 6.0) antigen retrieval (Vector, H-3300) at 95°C for 30 min and followed by blocking endogenous peroxidase activity with 3% H2O2 for 5 min. After three times of 5 min TBST washing, slides were blocked with blocking buffer (1.25% of goat serum) at room temperature for 1 h. Slides were applied with diluted primary antibody and incubated in a humidified chamber at 4°C overnight. After three 10 min TBST washes, slides were incubated with secondary antibody (Vector, PK-4001) for 30 min at room temperature the color of the antibody staining was revealed by peroxidase-based detection (Vector, SK4100) and the sections were counterstained with hematoxylin (Millipore Sigma, MHS1). Images were taken on a Leica Microscope with 20x objectives. The average of five images per tumor/tissue were scored based on the quantification of the % of cells negative, low (1+), medium (2+) or high (3+) immuno-intensity. Subsequently, histology/IHC score was calculated as H = [1X percentage of cells 1+] + [2 x (percentage of cells 2+)] + [3 x (percentage of cells 3+)]^72^.

### Puromycin incorporation SUnSET assay

#### In vitro

PCa cells were cultured under the specified conditions and pulsed with puromycin 10μg/ml for 10 mins in the culture medium. Cells were subsequently lysed, and immunoblotting was performed as previously described^55^.

#### In vivo

mice received an intraperitoneal injection of puromycin (40nmol/g bodyweight in 80uL) 20 mins prior to sacrifice. Xenograft tumor tissues were homogenized and lysed for immunoblotting as described above. Transgenic mouse tissues were fixed and embedded for immunofluorescence staining.

### TUNEL assay

Tissue slides were deparaffinized, rehydrated, antigen retrieval and goat serum blocking as described before. 50 μL of enzyme solution was added to 450 μL label solution to obtain 500 μL TUNEL reaction mixture. Tissue was incubated with the reaction mixture for 1 hr at 37°C, proceeded with three times of TBS washing and DAPI staining for 5 mins. Slides were covered with antifade mounting media (Vector Laboratories H-1900-2) and images were taken by EVOS FL microscope with 20x objectives.

### Immunofluorescence staining

Tissue slides were deparaffinized, rehydrated and performed antigen retrieval as described before, then followed by M.O.M. (Vector: MKB-2213-1) incubation for 1hr to block endogenous mouse immunoglobulins and then regular blocking using 5% goat serum for 2hrs at room temperature. Slides were incubated with primary antibodies (Puromycin, EMD Millipore MABE343, 1:400) overnight at 4°C. After three washes with TBS, slides were incubated with secondary antibodies (goat anti mouse 594 1:1000) for 1h at room temperature, then proceeded with three TBS washing and DAPI staining for 5mins. Slides were covered with antifade mounting media (Vector Laboratories H-1900-2) and images were taken by EVOS FL microscope with 20x objectives.

### LC-MS based metabolomic analysis

For *in vitro* cell line metabolites: three biological replicates of 2 million cells were used in each condition. Cells were washed twice with chilled PBS on ice and harvested with 1 ml of cold extraction solution (80% of methanol, 20% of water and 0.2 μM heavy standard mix Cambridge Isotope Laboratories, MSK-A2-1.2). Cell suspension was vortexed at 4°C and centrifuged at 13,000 rpm for 15min at 4°C. The supernatant was transferred to a pre-chilled cryovial without O-ring and kept at -80 C before metabolites mass spec. The pallets were air dried and were extracted for protein lysates. Protein concentrations were used for normalizing the metabolites amount. For tumor or animal tissue metabolites: 20mg tissue was snap frozen using liquid nitrogen and then homogenized using a pestle on dry ice. Metabolites were extracted with 500 μL of cold extraction solution by vortex and centrifuging at 13,000 rpm for 15 min at 4°C. The supernatant was transferred to a new 1.5 mL Eppendorf tube. The precipitate was re-extracted with 400 μL of cold extraction solution as described before. Then the combined supernatants were centrifuged and transferred to a pre-chilled cryovial without O-ring and kept at – 80°C before metabolites mass spec. The pallets were air dried and were extracted for protein lysates. Protein concentrations were used for normalizing the metabolites amount.

### Histone acid extraction and mass spectrometry sample preparation

Eight million cells were collected, washed with PBS, and centrifuged at 2500 rpm for 5 mins. Cell pellets were washed with 10 volumes of nuclei isolation buffer (NIB) and spun down. Nuclei were isolated by incubating the cells with 0.3% of Nonidet P-40 NIB for 5 mins. The nuclei were pelleted by centrifugation at 4°C and washed again with NIB alone. For acid histone extraction, 5 volumes of the original pellet of 0.4N H2SO4 were added to the isolated pellet and incubated for 3 hours at 4°C. Histones were centrifuged at 5200 rpm for 5 mins at 4°C and the histone containing supernatant was transferred to a new tube. One third of the supernatant volume of 100% trichloroacetic acid were added to the supernatant and incubated overnight at 4°C. Samples were centrifuged, washed ice-cold acetone with 0.1 HCl, centrifuged again and washed with 100% acetone. A final centrifugation was performed, and the histone pellets were air dried for 1 hour. Histones were resuspended in 50 μL of 100 mM NH4HCO3 and quantified. The obtained histones were used for immunoblotting. For histone profiling by liquid chromatography mass spectrometry, histones were derivatized with propionic anhydride by adjusting pH to 9 and incubated with a propionic anhydride with acetonitrile mixture at 37°C for 15 mins. Following the incubation period, samples were dried using nitrogen gas for 20-40 mins and resuspended in 100 mM NH4HCO3. The histones were then digested with trypsin overnight. The N-termini of the histone peptides was propionylated with Propionylation reagent and finally resuspended in water. Samples were desalted and ran by the Garcia Lab following their established protocol for histone post-translational modification studies.

### Histone mark immunoblotting

One microgram of histone protein was boiled in sample buffer, and 15-25 µL aliquots were separated by SDS-PAGE and transferred onto Polyvinylidene Difluoride membrane (GE Healthcare, IPVH00010). The membrane was blocked for one hour in blocking buffer [Tris buffered saline, 0.1% Tween (TBS-T), 5% nonfat dry milk] and then incubated overnight on a rocker at 4°C with the primary antibody. Following incubation, the blot was washed with TBS-T and incubated for 2 hours at room temperature with horseradish peroxidase(HRP)-conjugated secondary antibody. The blots were washed again, and signals were visualized using enhanced chemiluminescence system as per manufacturer’s protocol (GE Healthcare) or Kwik Quant Imager (Kindle Biosciences). Primary antibodies used in the immunoblotting assay are as follows: H3K27ac (Active Motif, 39133 – 1:1000), H3K23ac (Abcam, ab177275 – 1:1000), H3K36me3 (Abcam, ab194677 – 1:5000), H3K36me2 (Abcam, Ab 9049 – 1:5000), H3K27me3 (CST, C36B11 – 1:5000), H3K9me3 (Abcam, Ab 8898 – 1:5000), H3K4me3 (CST, C42D8 – 1:5000), H3K4me2 (CST, C64G9, 1:5000), H3K4me1 (Abcam, Ab 8895 – 1:5000), H4K20me2 (Abcam, Ab 9052 – 1:5000), phospho-H3 (EMD Millipore, 06-570 – 1:1000), and H3 (CST, D1H2 – 1:5000). Band intensities were quantified using ImageJ, histone marks were normalized to H3, and z-scores were calculated and plotted in heatmaps.

### 5mC Dot Blot

Genomic DNA was extracted using the QIAmp DNA Mini Kit. A working dilution sample of 10 ng/μL was prepared with TE Buffer (pH 8.0) and filtered 0.8N NaOH/20mM EDTA. Samples were vortexed and denatured at 95°C for 10 min in a thermocycler, followed by a 5 min incubation on ice. The N+ membrane and filter paper were hydrated in distilled water for 10 mins. The dot blot apparatus was assembled according to the Bio-Rad user manual. Samples were serially diluted by two-fold and loaded onto the dot blot apparatus. The membrane was air dried for 5 mins, exposed to UV light at 120,000 μJ/cm^2^, and block with 5% milk TBS-T at room temperature for 30 mins. One set of membranes was stained with methylene blue for DNA visualization. Another set of membranes were incubated with the 5mC antibodies from Diagenode (C15200081-100) or Millipore (MABE146) at a 1:1000 dilution overnight at 4°C. After secondary antibody incubation and subsequent washes, light intensity of the individual dots on the membrane was detected using the KwikQuant imaging system. Dot intensities were quantified using ImageJ, and fold changes were displayed in a bar graph.

### ChIP-Seq

For ChIP-seq experiments, VCaP and VCaP NNMT cells were grown in DMEM (high glucose, no glutamine, no methionine, no cystine; Gibco, 21013024) supplemented with dialyzed FBS (GeminiBio, 100-108). ChIP was performed using the iDeal ChIP-seq Kit for Histones (Diagenode, C01010059) according to manufacturer’s protocol. In brief, cells were cross-linked with 1% formaldehyde in culture medium for 8 minutes at room temperature. Cross-linking was quenched by adding 1/10 volume of 1.25 mol/L glycine and incubating for 5 mins at room temperature. Cells were lysed and chromatin was fragmented by sonication (Diagenode Bioruptor Pico, B01060010), resulting in an average chromatin fragment size of ∼250 bp. Chromatin equivalent to 5×10^6^ cells was isolated and incubated overnight at 4°C with 5 µg of the following antibodies, RNA Pol II, H3K27ac, H3K36me3, H3K36me2, H3K27me3, H3K4me3, H3K4me2, H3K4me1 and H3K9me3 Rabbit and Mouse IgG (Diagenode)]. During the ChIP reaction, drosophila chromatin equivalent to 2.5×10^5^ was added as a spike-in control for normalization. ChIP-seq libraries were prepared from the ChIP-enriched DNA samples using the NEBNext® Ultra™ II FS DNA Library Prep Kit for Illumina (E7805L). Briefly, 20 ng of ChIP-enriched DNA was converted to blunt-ended fragments. A single A-base was added to fragment ends, followed by ligation of sequencing adaptor. The adaptor-modified DNA fragments were enriched by PCR using unique NEB index primers. PCR products were purified using NEBNext Sample Purification beads (NEB, E6178S) and eluted with 1X TE buffer. Library size and quality were assessed using Agilent’s TapeStation system (G2991BA), and sequencing was performed on the Illumina NextSeq 2000 platform. Post-sequencing, read counts were normalized using a scaling factor calculated from the spike-in DNA across samples. This scaling factor was applied to adjust read counts of the experimental samples. DESeq2 ^74^ was employed to identify significant differences in chromatin binding events across all marks.

### RNA extraction

RNA was extracted using Qiagen’s RNeasy Mini kit (74104) according to manufacturer’s protocol. RNA integrity number was obtained by running the RNA samples in Agilent’s 4200 TapeStation system using the RNA ScreenTape Analysis kit (5067) according to manufacturer’s instructions.

### scRNA-seq tissue processing, library preparation, sequencing, and data analysis

Prostates from three *Pten^fl/fl^;Nnmt ^fl/fl^*, *Pten^-/-^*, and *Pten^-/-^;Nnmt^-/-^* mice dissected and processed to obtain single-cell suspensions using the Papain Dissociation System (Worthington Biochemical Corporation, LK003150). Cells were stained with 7-AAD and 7-AAD negative (live) cells were sorted at the Flow Cytometry Core at the Children’s Hospital of Philadelphia (CHOP). Sorted cells were submitted to the Center of Applied Genomics (CAG) for single cell processing using the 10X Genomics Chromium Next GEM Single Cell 3’ GEM, Library & Gel Bead Kit v3.1 (10X Genomics, PN-1000121). Libraries were sequenced on Illumina’s NovaSeq platform. The single-cell gene expression data was mapped to the mouse genome (mm10) using the Cell Ranger package (10x Genomics Cell Ranger 7.1.0). Cells with fewer than 200 expressed genes and greater than 20% mitochondrial reads were filtered out. Genes with less than 1 count were excluded from further analysis. Counts were normalized per cell to achieve one million reads per cell, and the top 1000 differentially expressed genes were selected for visualization using the SCANPY package^75^. Counts for these genes were scaled and log-transformed. Clustering was performed using the Louvain algorithm^76^ with default parameters. Principal Component Analysis (PCA) was then conducted on these 1000 genes, retaining the top 15 Principal Components (PCs). Visualization of the cells was achieved using the UMAP algorithm^77^, which was constructed with a neighborhood graph comprising 30 neighbors. Partition-based graph abstraction^78^ was utilized, with Louvain clusters serving as initial seeds for embedding. Cluster annotation was performed by identifying literature-defined markers for each cell type and annotating clusters based on marker expression. The markers used for cell lines were as follows: Endothelial: Cldn5, Vwf, Erg; Dendritic: Clec9a, Xcr1; B cells: Cd79a, Cd79b; T/NK cells: Cd3d, Cd3e, Cd3g, Cd4, Cd8a; Myeloid: Cd68, Aif1; Luminal: Krt8, Krt18, Cd24a, Psca, Cldn3, Aldh1a3, Epcam; Basal: Krt5, Krt14, Trp63; Seminal Vesicles: Pax2, Pate4, Calml3, M1 macrophage: Wnt2, Wnt6, Wnt10a, Rorb; Smooth Muscle cells: Acta2, Myh11; Glial: Sox10; Mesenchymal: CD44, Itgb1, Nt5e, Eng, Vim; Macrophage: Cd68, Cd14, Aif1; M2 macrophage – Rspo1, fgf10, Sult1e1. For Gene Set Enrichment Analysis (GSEA)^79^ only cells expressing luminal markers were included, yielding 1713 cells: 1046 from the *Pten^-/-^* condition and 667 from the *Pten^-/-^;Nnmt^-/-^* condition. Raw expression matrices for these cells were constructed, and phenotype based GSEA analysis was performed using the GSEApy package^80^ with the hallmark gene set.

### RNA-seq and ChIP-seq Analysis of Clinical data

Prostate cancer datasets from SU2C and TCGA were obtained from their respective repositories. For TCGA samples, patients were stratified into NNMT-high and NNMT-low groups based on an FPKM cutoff of ≥4.5, resulting in 325 high NNMT and 176 low NNMT patients. Similarly, for the SU2C dataset, an FPKM cutoff of 4 was used, yielding 64 high NNMT and 41 low NNMT patients. mRNA expression data from these groups were analyzed for hallmark pathway enrichment using GSEA^81^. For survival analysis, TCGA^43^ clinical data were matched to the corresponding expression dataset. Survival curves were generated using the ggsurvplot function from the R package survminer. For ChIP-seq, Tumor and matched normal samples from patients were downloaded from GEO accession GSE130408^82^. DeepTools^83^ was used to calculate matrices for H3K27me3 signal at all coding genes (±1 kb flanking regions) using the “scaled regions” method. These matrices were used to generate profile plots and heatmaps. Profile plots were created by averaging signal intensity across the three samples.

### Murine Prostate Tumor Xenograft Model

*In vivo* xenograft studies were performed in accordance with protocols approved by the Institutional Animal Care and Use Committee at the University of Pennsylvania and in compliance with all regulatory standards. Four- to 5-week-old male NOD/SCID mice were bred in house (Jackson Laboratory, 001303). For the prostate cancer cell xenograft experiments: 1-2 × 10^6^ cells suspended in 80 µL of RPMI-1640 with 50% Matrigel (BD Biosciences) were implanted subcutaneously into the dorsal flank of the mice. Tumor growth in all studies was recorded using digital calipers, and tumor volumes were estimated using the formula (π/6) (L × W2), where L = length and W = width of tumor. Body weight during the study was also monitored. At the end of the treatment regimen the mice were sacrificed and tumors extracted for downstream analysis.

For the methionine-restricted diet assays, mice were randomized into two groups and placed on either a methionine restricted diet containing 0.12% methionine (Research Diets, A11051301B) or a control diet with 0.86% methionine (Research Diets, A11051302B). The diets were initiated 14 days prior the tumor cell injection and maintained until the end of the assay. For the therapeutic assay, mice were randomized into four groups once tumors reached to 100mm^3^: group one was given control diet and vehicle oral gavage; group two was given methionine restrictive diet and vehicle oral gavage; group three was given control diet and oral gavage of Everolimus at 10 mg/kg body weight (50uL); group four was given methionine restricted diet with Everolimus treatment (10 mg/kg body weight,50uL). The treatment was administered for 24 to 28 days.

